# Phosphate-dependent nuclear export via a novel NES class recognized by exportin Msn5

**DOI:** 10.1101/2024.08.12.607649

**Authors:** Ho Yee Joyce Fung, Sanraj R. Mittal, Ashley B Niesman, Jenny Jiou, Binita Shakya, Takuya Yoshizawa, Ahmet E Cansizoglu, Michael P. Rout, Yuh Min Chook

## Abstract

Gene expression in response to environmental stimuli is dependent on nuclear localization of key signaling components, which can be tightly regulated by phosphorylation. This is exemplified by the phosphate-sensing transcription factor Pho4, which requires phosphorylation for nuclear export by the yeast exportin Msn5. Unlike the traditional hydrophobic nuclear export signal (NES) utilized by the Exportin-1/XPO1 system, cryogenic-electron microscopy structures reveal that Pho4 presents a novel, phosphorylated 35-residue NES that interacts with the concave surface of Msn5 through two Pho4 phospho-serines that align with two Msn5 basic patches, unveiling a previously unknown mechanism of phosphate-specific recognition. Furthermore, the discovery that unliganded Msn5 is autoinhibited explains the positive cooperativity of Pho4/Ran-binding and proposes a mechanism for Pho4’s release in the cytoplasm. These findings advance our understanding of the diversity of signals that drive nuclear export and how cargo phosphorylation is crucial in regulating nuclear transport and controlling cellular signaling pathways.

## Introduction

The regulation of gene expression in response to environmental stimuli hinges critically on the nuclear import and export of key components from signaling pathways like MAP kinase, TGF-β/SMAD, Hippo and others^1–3^. These proteins undergo dynamic changes in their localization between the nucleus and the cytoplasmic, controlled by phosphorylation, which either facilitates or inhibits interaction with nuclear transport receptors of the Karyopherin-β family, also known as importins, exportins and biportins^4^.

One such example is the *S. cerevisiae* (*Sc*) exportin Msn5 (also known as Kap142, homologous to human Exportin-5 or XPO5), which transports various transcription factors in response to nutrients and stress (reviewed in ^4^). Phosphorylation and dephosphorylation of these factors regulate their nuclear-cytoplasmic localization and transcription functions. Msn5 recognizes phosphorylated cargoes for nuclear export, such as the phosphate-sensing transcription factor *Sc* Pho4^5^. Pho4, regulated by phosphorylation from the Cyclin-CDK Pho80-Pho85 kinase complex, requires phosphorylation at specific sites for binding Msn5 and subsequent nuclear export^5–9^. However, until now, the precise Msn5-binding element, recognition mode and the mechanism linking phosphorylation to nuclear export have been unknown. Overall, understanding the broader mechanisms of nuclear export regulation by phosphorylation remains a significant challenge.

Exportins recognize their protein cargoes by binding to linear sequence elements known as nuclear export signals (NESs) or folded domains of the cargoes^4,10,11^. For example, the well-characterized Exportin-1 (XPO1 or CRM1) binds >1000 diverse functioning protein cargoes that carry the canonical-NES (cNES), a 8-15 residue long sequence rich in hydrophobic residues^12–16^. Most other exportins recognize folded domains in their cargoes; for example, profilin-actin by Exportin-6^17^, Importin-α by CSE1/CAS^18^, folded RNAs by XPOT and XPO5^19,20^, and various export cargo protein domains by biportins IPO13, XPO4 and XPO7^21–23^. The only other exportin known to bind IDRs of cargoes, via an embedded but until now uncharacterized NES, is Msn5^5^.

In this work, cryogenic electron microscopy (cryo-EM) analysis of Msn5 bound to Ran^GTP^ and phosphorylated Pho4 has revealed a 35-residue NES that binds in extended conformation to the concave surface of the flexible Msn5 solenoid. The Pho4 NES is anchored to Msn5 at two phospho-serine residues, which are surrounded by many small polar and hydrophobic residues that also make important interactions with the exportin. This study not only identifies a novel NES class for Msn5 distinct from the cNES recognized by XPO1 (previously the only characterized NES class) but also elucidates how a phosphorylated cargo is specifically recognized for nuclear export. These findings advance our understanding the mechanisms by which phosphorylation regulates nuclear export and their broader implications for a multitude of key cellular signaling pathways and protein transport mechanisms.

## Results and Discussion

### Mapping the Msn5-binding region of Pho4

The 312-residue Pho4 consists of a 240-residue IDR followed by a C-terminal helix-loop-helix domain^24,25^. The IDR contains a transactivation domain, an isoleucine-lysine (IK)-NLS that binds the importin Kap121, a putative oligomerization region, and two separate regions that recruit the Pho80-Pho85 kinase to phosphorylate five Pho4 serine residues within the IDR (Fig. 1a)^8,24,26,27^. We generated GST-fusion constructs of full length (FL) and truncated Pho4 to identify the minimal Pho4 region that binds Msn5. These GST-Pho4 constructs were phosphorylated *in vitro* with recombinant Pho80-Pho85 kinase. GST-pPho4_FL_, -pPho4_1-155_, -pPho4_1-200_ and -pPho4_1-255_ efficiently pulled down Msn5 in the presence of *Sc* Ran^GTP^ (residues 1-179, Q71L) but GST-Pho4_100-150_ did not pull down Msn5 (Fig. 1b and Extended Data Fig. 1a). We selected Pho4_1-200_, which contains both Pho80-binding regions and binds Msn5 like Pho4_FL_, for further studies.

**Fig. 1.**
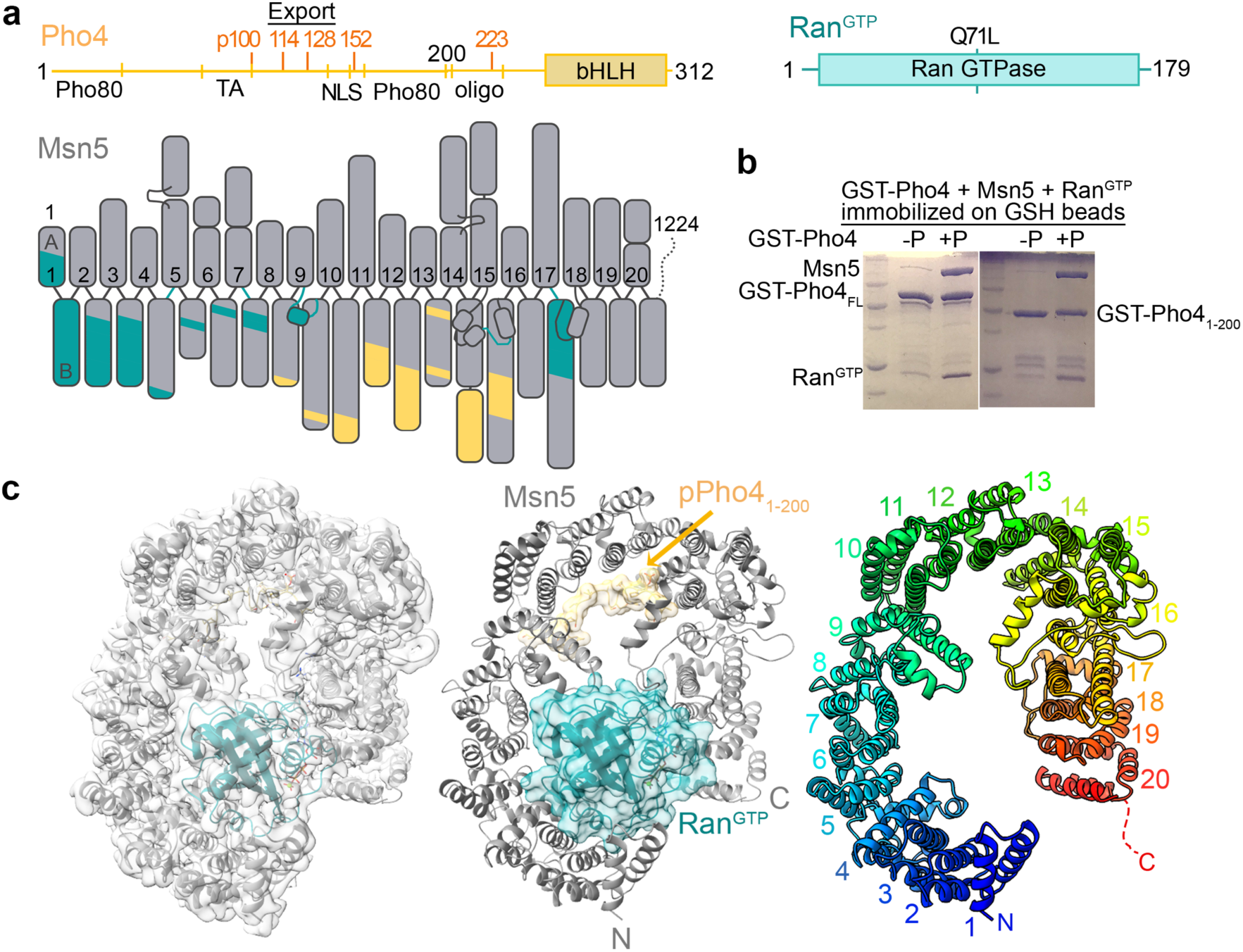
Cryo-EM structure of Msn5-Ran^GTP^-pPho4. **(a)** Schematic of Pho4, Ran^GTP^ and Msn5. Pho4 has two Pho80-interacting regions (Pho80), a transactivation domain (TA), an IK-NLS, an oligomerization region (oligo) and five Pho85 phosphorylation sites (in orange) in its IDR, and a C-terminal basic helix-loop-helix domain (bHLH). Each grey box in the Msn5 schematic represents a helix scaled according to its length (small helices in h9loop, h15loop and h17-h18loop are also shown). HEAT repeats are numbered and the regions that contact Ran^GTP^ or pPho4 are cyan or yellow, respectively. **(b)** Immobilized GST-Pho4_FL_ and -Pho4_1-200_, *in vitro* phosphorylated (+P) or unphosphorylated (-P), were mixed with Msn5 and excess Ran^GTP^ followed by extensive washing. Bound proteins were visualized by Coomassie stained SDS-PAGE. See also **Extended Data Fig.1**. **(c)** Left: The final map (transparent grey) overlayed with the Msn5 (grey)-Ran^GTP^(cyan)-pPho4_1-200_(wheat) structure in cartoon (also shown alone in the center with Ran^GTP^/pPho4 in transparent surfaces). Right: Msn5 shown alone in rainbow colors with HEAT repeats labeled. See also **Extended Data Fig. 2-4**.

Fluorescence polarization (FP) analysis showed high affinity binding of wild type or WT mNeonGreen(mNeonG)-pPho4_3-200_ to WT Msn5 (dissociation constants or K_D_ 25[14,43] nM; 95% confidence interval in brackets) (Extended Data Fig. 1b and c). The interaction of mNeonG-pPho4_FL_ with WT Msn5 showed 2-site binding, with K_D_s of 10[U,90] nM (U = undetermined) and 400[200,2200] nM, consistent with a dimeric Pho4_FL_ binding with high affinity to Msn5 (Extended Data Fig. 1b and c).

### Cryo-EM maps of pPho4_1-200_- and pPho4_FL_-bound Msn5 reveal Msn5 flexibility

We purified ternary complexes of Msn5-Ran^GTP^-pPho4_1-200_ and Msn5-Ran^GTP^-pPho4_FL_ for cryo-EM structure determination. We solved a 3.0 Å resolution structure of the former and obtained a 4.9 Å resolution map of the latter (Fig. 1c, Table 1, Extended Data Fig. 2). 3D reconstruction of Msn5-Ran^GTP^-pPho4_1-200_ particles showed the 20 HEAT repeats (h1-h20; each with antiparallel A and B helices) of the horseshoe-shaped Msn5 solenoid, with Ran^GTP^ bound to its N-terminal repeats. 3D variability analysis revealed conformational variability of the Msn5 solenoid, prompting us to split the particles into six classes (Extended Data Movie 1). The resulting maps revealed Msn5 conformations that vary in the extent of solenoid opening (Extended Data Fig. 2-3). Similarly, reconstruction of the Msn5-Ran^GTP^-pPho4_FL_ particles yielded maps that show different extent of Msn5 solenoid opening (Extended Data Fig. 2-3).

**Table 1.** Cryo-EM data, map and model statistics of Msn5-Ran^GTP^-pPho4_1-200_ and unliganded Msn5.

|  | **Msn5-Ran<sup>GTP</sup>-pPho4<sub>1-200</sub>** | **Msn5** |
| --- | --- | --- |
| **Data collection and processing** |  |  |
| Magnification | 105,000 | 81,000 |
| Voltage (kV) | 300 | 300 |
| Electron exposure (e<sup>-</sup>/Å<sup>2</sup>) | 50 | 50 |
| Defocus range (μm) | -0.8 to -2.5 | -0.8 to -2.5 |
| Pixel size (Å) | 0.83 | 1.02885 |
| Symmetry imposed | C1 | C1 |
| Initial particle images after 1<sup>st</sup> round of clean-up (no.) | 1,574,329 | 4,135,768 |
| Final particle images (no.) | 102,030 | 1,048,421 |
| Map resolution (Å) | 3.08 | 3.39 |
| FSC threshold | 0.143 | 0.143 |
| B factor (Å<sup>2</sup>) | 89.9 | 174.5 |
| **Refinement** |  |  |
| Initial model used | Msn5 and Ran<sup>GTP</sup> from 3m1l | SWISS model of 5YU7 |
| Model resolution (Å) | 2.9/3.0/3.2 | 3.3/3.4/3.6 |
| FSC threshold | 0/0.143/0.5 | 0/0.143/0.5 |
| Model composition |  |  |
| Nonhydrogen atoms | 11123 | 9754 |
| Protein residues | 1354 | 1189 |
| Ligand | 2 (GTP, 1 Mg) | 0 |
| R.m.s. deviations |  |  |
| Bond lengths (Å) | 0.003 | 0.002 |
| Bond angles (°) | 0.610 | 0.481 |
| **Validation** |  |  |
| MolProbity score | 1.33 | 1.53 |
| Clashscore | 5.43 | 5.52 |
| Poor rotamers (%) | 0.72 | 0.09 |
| Ramachandran plot |  |  |
| Favored (%) | 97.84 | 96.45 |
| Allowed (%) | 2.16 | 3.55 |
| Disallowed (%) | 0.00 | 0.00 |
| PDB/ EMDB code | 9D45/EMD-46549 | 9D43/EMD-46548 |

Map features for the bound pPho4 in all the Msn5-Ran^GTP^-pPho4_1-200_ and Msn5-Ran^GTP^-pPho4_FL_ maps can be observed in the concave surface of Msn5, but the extent and continuity vary across the maps (Extended Data Fig. 4). The most compact states of both complexes are very similar, have the strongest and most continuous Pho4 feature, and are described below. The variable Msn5 solenoid and Pho4 conformations in the different maps, all from a single cryo-EM sample, suggest dynamic Pho4-Msn5 interactions with the flexible Pho4 polypeptide adapting to bind Msn5 solenoids of variable curvatures.

### A 35-residue NES of Pho4 binds the central concave surface of Msn5

We solved the structure of the most compact state of the Msn5-Ran^GTP^-pPho4_1-200_ complex (Fig. 1c). We built the h1-h19 repeats with confidence and placed h20 by alignment into less well-defined density (Extended Data Fig. 2b). The 20 HEAT repeats of Msn5 include a few unusually large HEAT repeats (h14, h15 and h17) and long loops (h9loop, h15loop and h17-h18loop) (Fig. 1a and c). The mode of Ran-binding is like that of other Kaps^4^: Msn5 h1-h3 bind the Ran^GTP^ switch II, h4-h9 and h9loop bind the Ran^GTP^ basic patch, while h15loop and h17 bind Ran switch I (Extended Data Fig. 5).

Strong and continuous map features adjacent to the concave surface of Msn5 h8-h15 allowed confident modeling of 23 Pho4 residues (Fig. 2a– c and Extended Data Fig. 2c). The bound Pho4 residues 112-134 form a large and sprawling 1256.9Å^2^ interface with Msn5. This Pho4 segment is likely the persistently bound core portion of the Pho4 NES. The same Pho4 segment is resolved in the most compact Msn5-Ran^GTP^-pPho4_FL_ map, with additional map features beyond the N-terminus of Pho4 L112, adjacent to a basic patch on Msn5 h16-h18 (Fig. 2d). We estimate that Pho4 residues 100-111 occupy this density, as mutating the electronegative ^102^ATTATI^107^ to GKKGKK decreased Msn5 affinity by 6-fold (K_D_ = 110[70,150] nM for 102-107_GKK_ *vs* 20[12,31] nM for WT; Fig. 2e). Altogether, cryo-EM results reveal a Pho4 NES spanning residues 100-134 that zigzags along the B helices of Msn5 h8-h18 in an overall opposite direction to the solenoid (Fig. 2b).

**Fig. 2.**
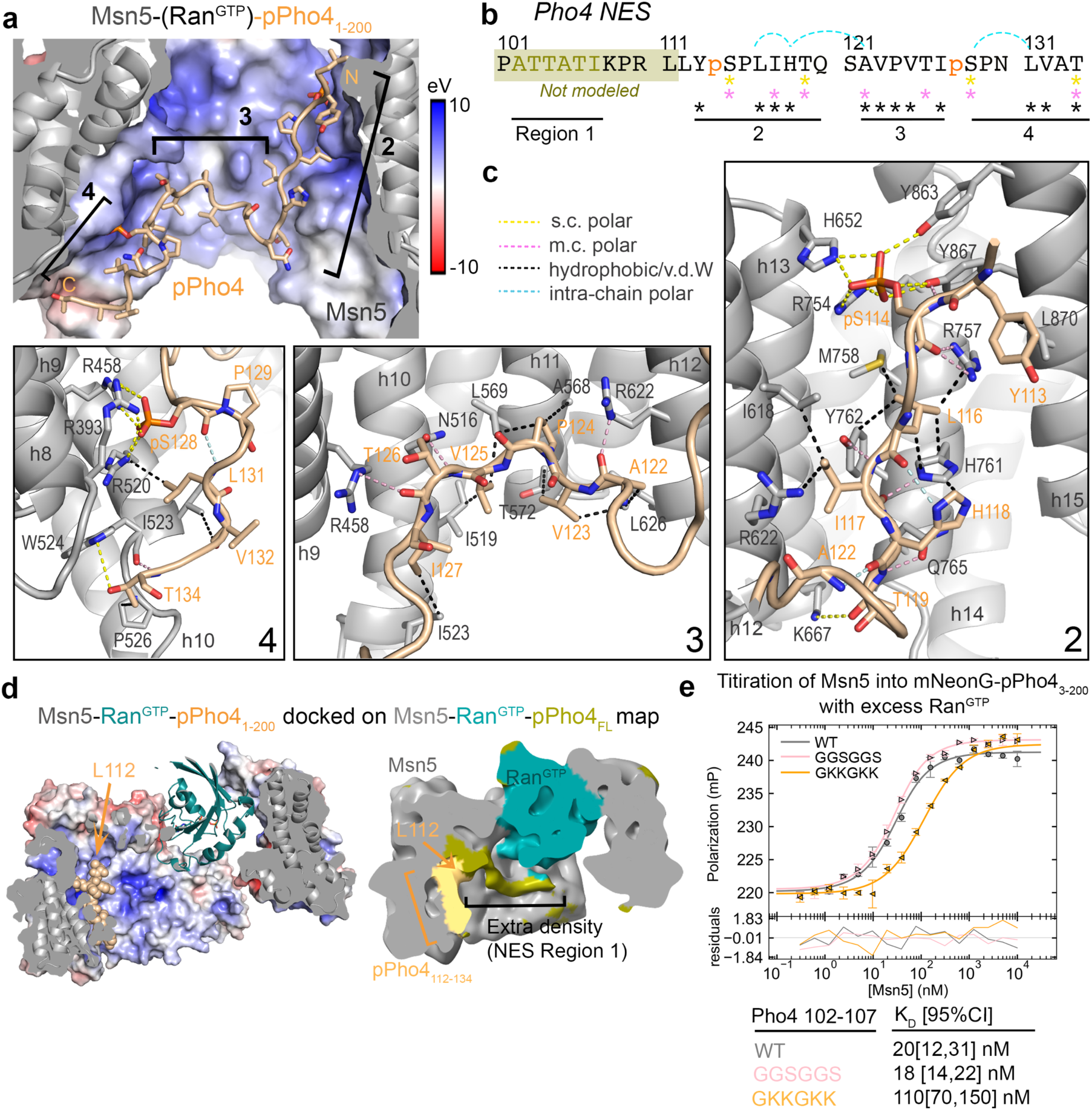
pPho4 NES binds to the central concave surface of Msn5. **(a)** Msn5 bound to pPho4_112-134_ (wheat cartoon; regions 2 – 4 labeled). Msn5 electrostatic surface potential (red to blue: −10 to 10 eV; same in **d**). See also **Extended Data Fig. 2c**. **(b)** The pPho4 NES sequence (regions 1-4 labeled) with Msn5 contacts represented by yellow, pink, and black asterisks - side chain polar, main chain polar and hydrophobic contacts respectively, blue dotted lines - intra-chain polar. Residues in olive box were not modeled. **(c)** Details of pPho4 NES regions 2 – 4, with contacts shown as dotted lines in same color scheme as **b**. **(d)** State 1 map of Msn5-Ran^GTP^-pPho4_FL_ (right) colored as in the docked Msn5-Ran^GTP^-pPho4_1-200_ structure (left, in the same orientation, colored as in **a**, pPho4 in spheres). Extra continuous density N-terminal of the pPho4_112-134_, likely occupied by Pho4 NES region 1, is colored in olive. **(e)** FP titration of Msn5 into mNeonG-Pho4_3-200_ with excess Ran^GTP^. Residues 102-107 (NES region 1) were mutated to GGSGGS or GKKGKK. Data points are mean ± s.d. from triplicates, line represent 1-site binding and fitting residuals are plotted below.

### The four regions of the Msn5-bound Pho4 NES

We divided the 35-residue Pho4 NES into four regions based on the locations of the three 90° turns of the zig-zagging chain (Fig. 2a and b). From the N-terminus, ^100^**pS**PATTATIKPRL^111^ (NES region 1) binds dynamically to the basic patch at Msn5 h16-h18 (Fig. 2d). The peptide then makes a 90° turn at Pho4 L112 and ^112^LY**pS**PLIHTQ^120^ (NES region 2) binds in a shallow basic/hydrophobic Msn5 groove formed by HEAT repeats h12-h15 (Fig. 2c, right panel). The Pho4 chain makes another ∼90° turn at ^120^QS^121^ placing ^122^AVPVTI^127^ (NES region 3) over a hydrophobic patch formed by Msn5 helices h10B, h11B and h12B (Fig. 2c, middle panel). A last ∼90° turn at pS128 places the ^128^**pS**PNLVAT^134^ region (NES region 4) approximately parallel to the adjacent basic and acidic surfaces of Msn5 h8-h10 (Fig. 2c, left panel).

It is obvious from the structure that phosphorylated Pho4 100-134 is bound to Msn5, but this Pho4 segment alone is insufficient for Msn5 binding because it is missing the kinase binding sites and therefore cannot be phosphorylated efficiently. Synthetic phosphopeptides were unsuitable for binding studies because of aggregation. Interestingly, phosphomimic mutations do not mimic Pho4 phosphorylation: Pho4_1-200_ S114/128_DD_ and Pho4_1-200_ S114/128_EE_ bind Msn5 weakly, like unphosphorylated Pho4_1-200_, with K_D_s ∼2-3 µM (Extended Data Fig. 6a).

### Msn5 interactions involving Pho4 pS114 and pS128

Phospho-serines pS114 and pS128 were previously reported to be critical for Msn5-mediated nuclear export^5^. The phosphate moiety of Pho4 pS114 interacts with side chains of Msn5 residues H652, R754, Y863 and Y867 within repeats h13-h15 while pS128 interacts with three R393, R458 and R520 side chains within Msn5 h8-h10 (Fig. 3a). Mutation of Pho4 S114 or S128 to alanine decreased Msn5 binding affinity by 6- and 3-fold respectively (K_D_ = 250[170,370] nM for S114_A_ and120[90,160] nM for S128_A_ *vs* 44[26,71] nM for WT; Fig. 3b, left panel). Mutating both serines (S114/128_AA_) decreased affinity 23-fold (K_D_ 1[0.7,1.4] µM), which is ∼2-fold tighter than unphosphorylated pPho4 (K_D_ 2.4[1.9,3.2] µM) (Fig. 3b, left panel and Extended Data Fig. 6a). This led us to examine the two remaining phosphoserines pS100 and pS152 that are not observed in the structure. Mutation of S100/152 to alanines decreased affinity by 2-fold (K_D_ = 46[38,56] nM for S100/152_AA_ *vs* 20[12,31] nM for WT; Extended Data Fig. 6b). Altogether, these mutagenesis/binding affinity studies show that phosphorylation of Pho4 S114 and S128 is key for Msn5-binding while phosphorylation of S100 and S152 contributes minimally.

**Fig. 3.**
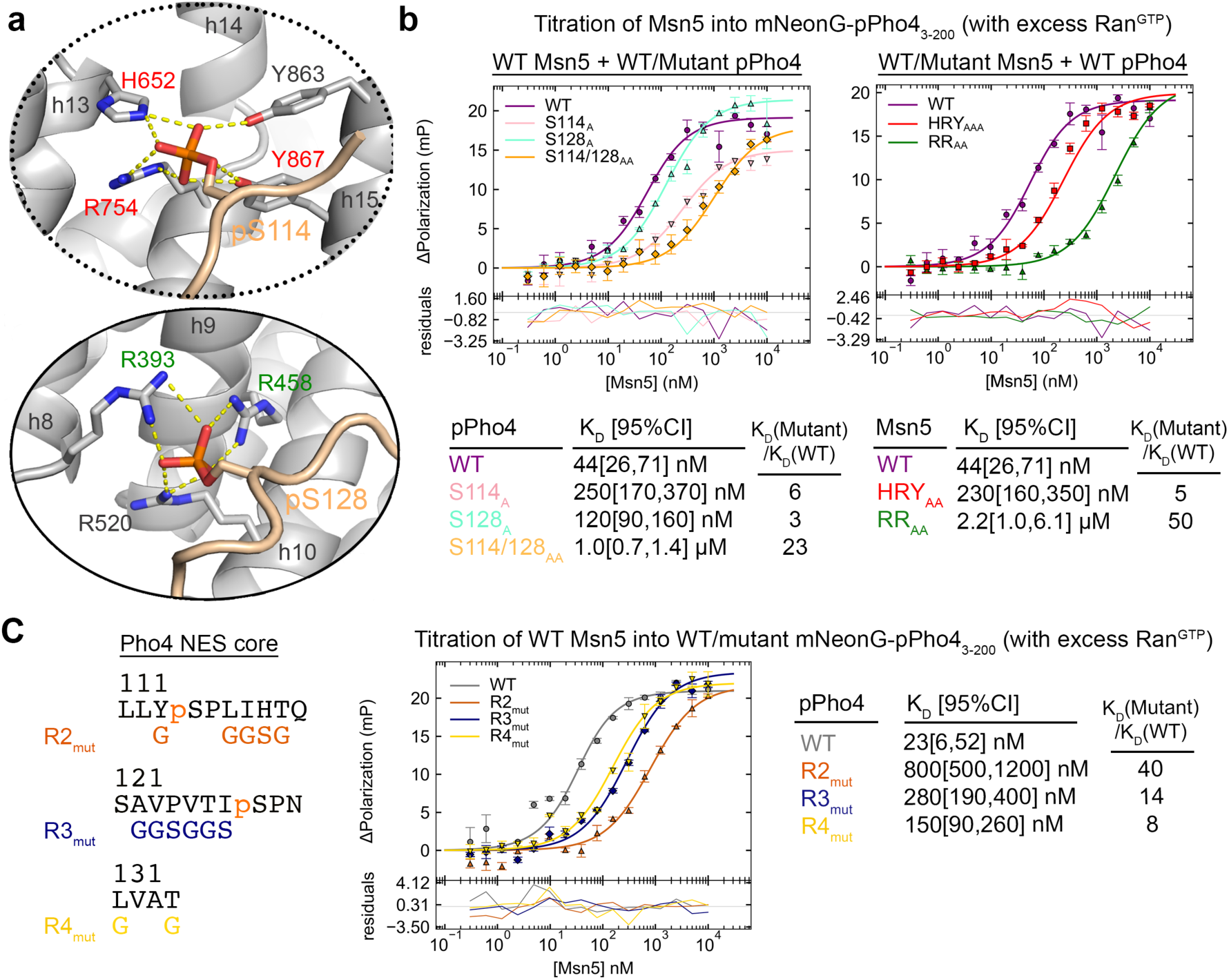
Pho4 phosphates and hydrophobic/polar side chains contribute to high affinity pPho4-Msn5 interactions. **(a)** Msn5 residues that contact pS114 and pS128. Residues mutated in Msn5 HRY_AAA_ are labeled in red and those in Msn5 RR_AA_ are in green. **(b)** Left: FP assays of WT Msn5 binding to mNeonG-pPho4_3-200_ (WT and S114/S128 mutants) in the presence of excess Ran^GTP^. Right: WT, HRY_AAA_ and RR_AA_ Msn5 binding to WT mNeonG-pPho4_3-200_ with excess Ran^GTP^. Data points represent mean ± s.d. of triplicate measurements. Lines represent 1-site binding fits and residuals are plotted below. Dissociations constants (K_D_) are shown with 95% confidence interval in brackets. **(c)** As in left panel of **b**, but with pPho4 NES mutated in regions 2-4, as indicated on the left. **See also Extended Data Fig. 6**.

We also mutated the pS114-binding Msn5 residues (H652, R754, Y867 or the Msn5 HRY site; HRY_AAA_) and the pS128-binding Msn5 residues (R393 and R458 or the Msn5 RR site; RR_AA_). Both mutants bound pPho4 weaker: Msn5 HRY_AAA_ bound 5-fold weaker (K_D_ 230[160,350 nM) and Msn5 RR_AA_ bound 50-fold weaker (K_D_ 2.2[1.0,6.1] µM) (Fig. 3b, right panel). This show that while both Msn5 phosphate-binding hotspots HRY and RR are important for pPho4 binding, RR is the stronger of the two. Comparison of these Msn5 mutants with the respective pPho4 S114_A_ and S128_A_ mutants and the binding studies of Msn5 mutants binding to Pho4 mutants are described in supplementary text.

In summary, the cryo-EM structure shows that Pho4 pS114 binds at the Msn5 HRY site, and pS128 binds at the Msn5 RR site. Mutagenesis/affinity analyses validate these interactions and show that at least one of the Pho4 phosphoserines is needed for sub-micromolar affinity pPho4-Msn5 binding. Interactions at the Msn5 RR site contribute more binding energy than the Msn5 HRY site.

### Transcription factor cargoes binding to Msn5 HRY and RR sites regulates expression of downstream genes

To examine the functional relevance of these *in vitro* findings *in vivo*, a Msn5 knockout yeast strain was acquired into which either an empty vector control, a WT Msn5, or a Msn5 construct with both phosphate binding HRY and RR hotspots mutated (HRYRR_AAAAA_) was transformed. These strains were grown to mid-log growth phase and their total RNA was extracted and subjected to mRNA sequencing (Fig. 4). Gene expression changes comparing the empty vector control strain to the WT Msn5 strain as well the HRYRR_AAAAA_ mutant strain to the WT Msn5 strain were determined and plotted. Our *in vitro* analyses indicated that the mutant strain should be binding incompetent, so export of Pho4 and other Msn5 transcription factor cargoes that bind at the Msn5 HRY and RR sites should be nearly abolished. As such, in both the empty vector control and the mutant strain, transcription factor cargoes should remain in the nucleus and the resulting expression of genes under the control of these transcription factors should increase.

**Fig. 4.**
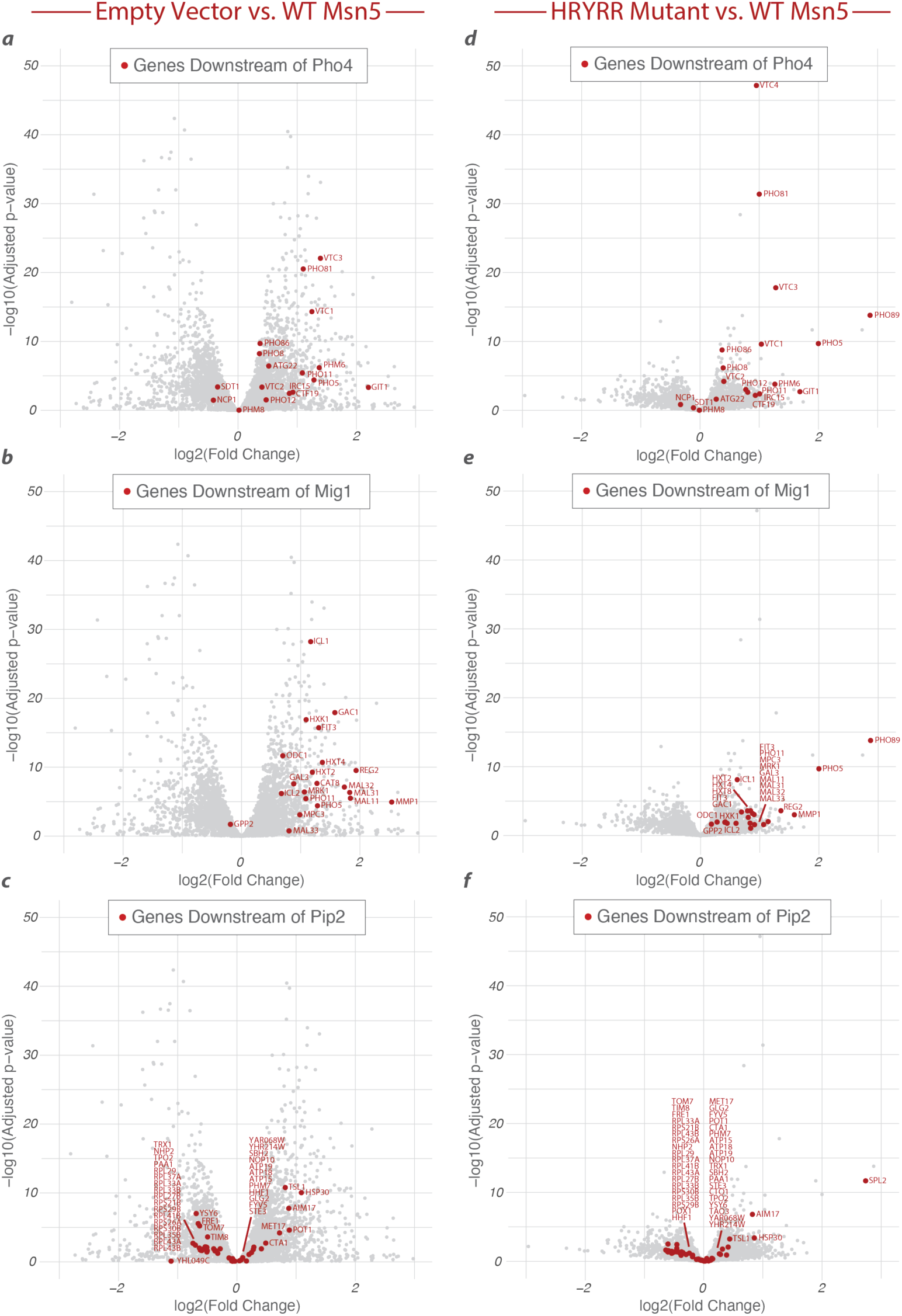
Genes downstream of Msn5 transcription factor cargoes show similar pattern of expression changes in Msn5 knockout and HRYRR_AAAAA_ mutant strains. **(a-c)** Volcano plots comparing Empty Vector (Msn5 KO) gene expression to Msn5 WT gene expression. **(d-f)** Volcano plots comparing Msn5 HRYRR_AAAAA_ mutant strain gene expression to WT Msn5 gene expression. Genes downstream of the transcription factors Pho4 (**a,d**), Mig1 (**b,e**), and Pip2 (**c,f**) are highlighted in red.

Beginning with the Pho4 transcription factor (Fig. 4a and d), genes downstream show a general trend towards overexpression as compared to the WT Msn5 strain in both the Empty Vector and HRYRR_AAAAA_ mutant conditions. When this same analysis is applied to the Mig1 transcription factor (Fig. 4b and e), another cargo of Msn5, we see the same general trend towards overexpression in both the empty vector and mutant strains. Finally, as a control, we plotted the genes downstream of Pip2 (Fig. 4c and f), a transcription factor that does not bind Msn5, and there is no discernible trend in either condition.

The expression of the binding incompetent double mutant results in a similar trend of increased expression of genes under the control of its cognate cargoes as does expressing an empty vector in an Msn5 knockout background. We have also shown that these effects are not generalized across all transcription factors but limited specifically to those that are directly exported from the nucleus by Msn5. In summary, this RNAseq analysis has validated the critical functional importance of the HRY and RR binding pockets in yeast and has also highlighted an important function of Msn5 in regulating gene expression.

In keeping with previous results showing that overexpression of Msn5 results in a slower growth phenotype^28–30^, equivalent overexpression of both the WT and HRYRR_AAAAA_ mutant proteins were equally deleterious to cell growth on equivalent over expression (Extended Data Fig. 7).

### Hydrophobic and polar side chains of pPho4 NES contribute to Msn5-binding

Fourteen other non-phospho-serine residues within the Pho4 NES core also make extensive interactions with Msn5 (Fig. 2b and c). The side chains of NES region 2 (^113^YpSPLIHTQ^120^) make numerous hydrophobic interactions with side chains of the B helices of Msn5 h12-h15, while the Pho4 main chain makes many polar interactions with Msn5 (Fig. 2c, right panel). Interacting Msn5 residues here are mostly hydrophobic or basic. Additionally, intramolecular Pho4 interactions, such as between the main chain carbonyl of H118 with the main chain amide of A122 likely stabilizes the 90° turn of the Pho4 chain. Mutating all the non-phospho-serine Msn5-interacting residues in region 2 (^113^YpSPLIHT^119^ to GpSPGGSG; R2_mut_) decreased Msn5-binding by 40-fold (K_D_ 800[500,1200] nM; Fig. 3c), underscoring the substantial contributions of these non-phospho-serine residues to Msn5 binding (Fig. 3b).

Every Pho4 side chain in NES region 3, ^122^AVPVTI^127^, contacts Msn5 (Fig. 2c, middle panel). The small hydrophobic side chains of Pho4 residues 122-125 make hydrophobic interactions with Msn5 side chains in h10-h12, and both ends of the NES region 3 main chain make polar interactions with Msn5 arginine side chains. Intramolecular Pho4 interactions are also present, with the V123, V125 and I127 side chains lining up on one side of the Pho4 chain, contacting each other. Mutating ^122^AVPVTI^127^ to GGSGGS (R3_mut_) decreased affinity 14-fold (K_D_ 280[190,400] nM; Fig. 3c), supporting the importance of these side chains in binding Msn5.

Pho4 NES region 4 (^128^pSPNLVAT^134^) contacts the B helices of Msn5 helices h8-h10 (Fig. 2c, left panel). ^128^pSPNL^131^ forms a turn held by intramolecular hydrogen bond-like contacts, which position pS128 and L131 onto a basic patch at the Msn5 RR site. Residues before and after the turn also interact with Msn5 residues. Mutations of the only non-phospho-serine side chains here that contact Msn5, Pho4 L131A/T134A (R4_mut_), decreased affinity by 8-fold (K_D_ 150[90,260] nM; Fig. 3c), supporting their importance in Msn5-binding.

In summary, many hydrophobic and small polar side chains in each of Pho4 NES regions 2, 3 and 4 contribute substantially to total binding energy. They are as important as phosphoserines pS114 and pS128 for high-affinity Msn5-binding. Importantly, though, many Pho4-Msn5 contacts also involve the Pho4 main chain.

### Characteristics of the Pho4 NES and its binding site on Msn5

Structural analysis shows a 35-residue Pho4 NES that zigzags across a long contiguous interface on the concave and hydrophobic/basic surface that spans HEAT repeats h8-h18 of Msn5. The Pho4 NES includes the dynamic N-terminal region 1 followed by the persistently bound core of regions 2 – 4. Sixteen of the 23 Pho4 NES core residues contact Msn5. These include two critical phosphoserines pS114 and pS128 positioned 30 Å apart into two binding hotspots of Msn5 (HRY and RR sites), and many hydrophobic and small polar side chains that are also important for binding. The Pho4 NES also uses many main chain atoms across the NES to bind Msn5. Furthermore, intramolecular hydrogen-bond like contacts within the NES enable the chain to make the three sharp turns to stay close to the curved Msn5 interface. ConSURF analysis of 101 fungal Msn5 homologs show that 79% (37 of 43) of the residues that make up the Pho4-binding interface are conserved, consistent with their roles in binding Pho4 and likely other Msn5 cargoes (Extended Data Fig. 8).

Based on the Pho4 NES and Msn5 features above, we propose that the following NES characteristics may be generally important for binding Msn5: 1) Location in an IDR. 2) Two potential phosphorylation sites spaced 10 residues or more to span the 30 Å between the Msn5 HRY and RR sites. 3) Enrichment in aliphatic hydrophobic and small polar side chains. 4) Few or no basic residues since the Pho4-binding site on Msn5 is basic. We examined sequences within the regions in 16 putative Msn5 cargoes that were reported to be important for nuclear export. Sequences that seem to fit the criteria noted, found in eight of the proteins, are listed in Extended Data Table 2. However, the lack of knowledge about kinases that control their nuclear/cytoplasmic localization and phosphorylated sites that drive nuclear export severely hampers testing of NESs that might bind Msn5.

The Pho4 NES binding site that spans HEAT repeats h8-h15 of Msn5 is large and open, and the Pho4 NES binds only two of several basic patches on this surface. We therefore predict that Msn5-binding NESs might be very diverse in sequence and structure, with phospho-serines or -threonines potentially binding to different combinations of basic Msn5 patches. The extent of the structural and sequence diversity of the NESs that bind at repeats h8-h18 of Msn5 will only be evident with additional cargo-bound structures.

### Cargo recognition by Msn5 *vs* XPO5

Msn5 and its human homolog XPO5 share 21% sequence identity. Alignment of the cargo-bound exportin structures using the DALI server^31^ shows similar HEAT repeat organization and architecture (Extended Data Fig. 9). Many yeast and human Kap homologs share conserved cargo specificities, but Msn5 is known to export phosphorylated proteins while XPO5 exports pre-miRNAs^32–36^. Both XPO5 and Msn5 were reported to export tRNAs and XPO5 was reported to also export dsRNA-binding proteins. It is unclear if XPO5 binds those proteins directly or via their common RNA ligands^37–45^.

Alignment of the Msn5 and XPO5 structures shows that both exportins use their concave surfaces at h8-h19 to bind Pho4 and pre-miRNA, respectively (Fig. 5a). However, the binding sites overlap only partially. Furthermore, only 11 of 27/41 Msn5/XPO5 residues that bind Pho4/pre-miRNA are in common and only three Msn5 residues that contact Pho4 are conserved in XPO5 (Extended Data Fig. 9). All three residues, R393, R458 and R520, are part of the dominant phosphate-binding hotspot that binds Pho4 pS128, the Msn5 RR site, while the equivalent arginine residues in XPO5 (R380, R448 and R519) interact with the dsRNA stem (Fig. 5b). While the Msn5 RR site is conserved in XPO5, the Msn5 HRY site is not. Overall, only ∼30% of the residues that contact Pho4 or pre-miRNA are similar/identical in Msn5/XPO5. Partial conservation of the cargo binding mode in Msn5 and XPO5 suggests a partial conservation of their protein cargo/NES specificities.

**Fig. 5.**
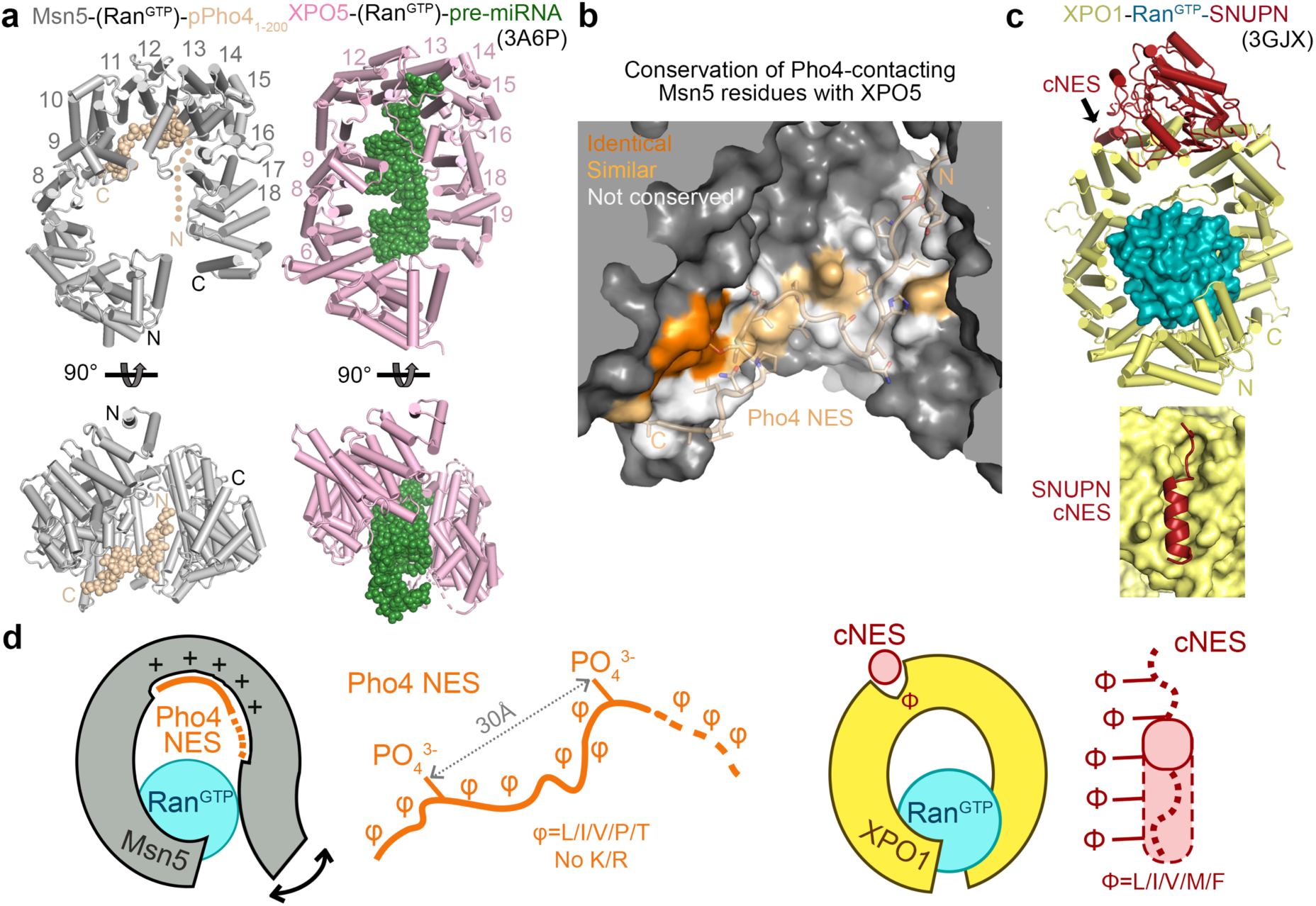
Comparison of cargo-bound Msn5 with XPO5 and XPO1. **(a)** Alignment of Ran^GTP^ in the Ran/cargo-bound structures Msn5 (grey) and XPO5 (pink; 3A6P). Ran^GTP^ is omitted for clear view of the cargoes, which are drawn as spheres - pPho4 in wheat and pre-miRNA in green. HEAT repeats that contact the cargoes are labeled with their numbers. **(b)** Conservation of Msn5 residues that contact pPho4 (via DALI structural alignment of unliganded Msn5 and XPO5 structures). The Msn5 surface and pPho4 cartoon are shown. Msn5 residues that contact pPho4 are colored orange if identical, light orange if similar and white if not conserved with the homologous XPO5 residues. See also **Extended Data. Fig. 9**. **(c)** Top: XPO1 (yellow)-Ran^GTP^ (teal)-SNUPN (red) (3GJX) oriented as in **a**. Bottom: View of the SNUPN cNES in the NES binding groove of XPO1 (viewing in the direction of arrow in the top panel). **(d)** Cartoon summarizing Msn5-Pho4 NES (left) *vs*. XPO1-cNES (right) recognition. The Pho4 NES is an extended 35-amino acids-long chain with two phospho-serines spaced 14 residues (30 Å) apart and many small hydrophobic and polar residues. It is depleted of basic side chains as it interacts with the highly basic and flexible concave surface of Msn5. In contrast, the cNES is short (8-15 residues long), with 4-5 hydrophobic side chains that bind in a hydrophobic groove located on the convex surface of XPO1. The groove holds the same conformation as it binds structurally variable cNESs (dashed lines depict variability, solid line shows the structurally conserved single turn of helix).

### The Pho4 NES bears no resemblance to the cNES that binds XPO1

The NES of Pho4 is strikingly different from the cNESs that bind XPO1 (Fig. 1c, 2a, 5c and d) in several ways: 1) The much longer 35-residue Pho4 NES core adopts an extended conformation when bound to Msn5, while the 8-15 residue cNESs adopt diverse helix-loop structures when bound to XPO1^12–16^. 2) The Pho4 NES binds to a large, flexible and basic binding site on the concave side of the Msn5 solenoid, whereas cNESs bind to an invariant hydrophobic groove on the convex side of the XPO1 solenoid. 3) The binding energy for Msn5-Pho4 interactions is distributed across 4 regions spanning 35 amino acids of the NES and involves two phosphoserine anchors. Such energetically distributed interactions are also found in PY-NLSs that use 3-4 NLS epitopes to bind the importin TNPO1/Kapβ2^46,47^. In contrast, cNES-XPO1 interactions are anchored by 4-5 hydrophobic side chains within a compact ∼8-15 residues long peptide^12,14,48^. 4) Msn5 binds pPho4 with low nanomolar affinity and Msn5/Pho4 mutants that bind at low micromolar affinities are no longer active for nuclear export^5^. In contrast, the cNES-XPO1 affinity range for active nuclear export is large, spanning low nanomolar to tens of micromolar^49^. The narrow affinity range for active nuclear export by Msn5 may be important for tight regulation of nuclear export in response to signaling from the environment.

### Msn5 autoinhibition, pPho4/Ran^GTP^ positive cooperativity and Pho4 release

We also solved the cryo-EM structure of unliganded Msn5 to 3.4 Å resolution (Fig. 6a, Table 1 and Extended Data Fig. 10). Unliganded Msn5 particles show less flexibility than Pho4-bound Msn5 (Extended Data Fig. 10a). Unliganded Msn5 is ring-shaped and more compact than the most compact pPho4/Ran^GTP^-bound Msn5, much like its human homolog XPO5 (5YU7 and 3A6P)^20,50^ (Fig. 6a, overlays in Extended Data Fig. 11a). The closed Msn5 ring is stabilized by extensive intramolecular interactions (Fig. 6a and Extended Data Fig. 10d).

**Fig. 6.**
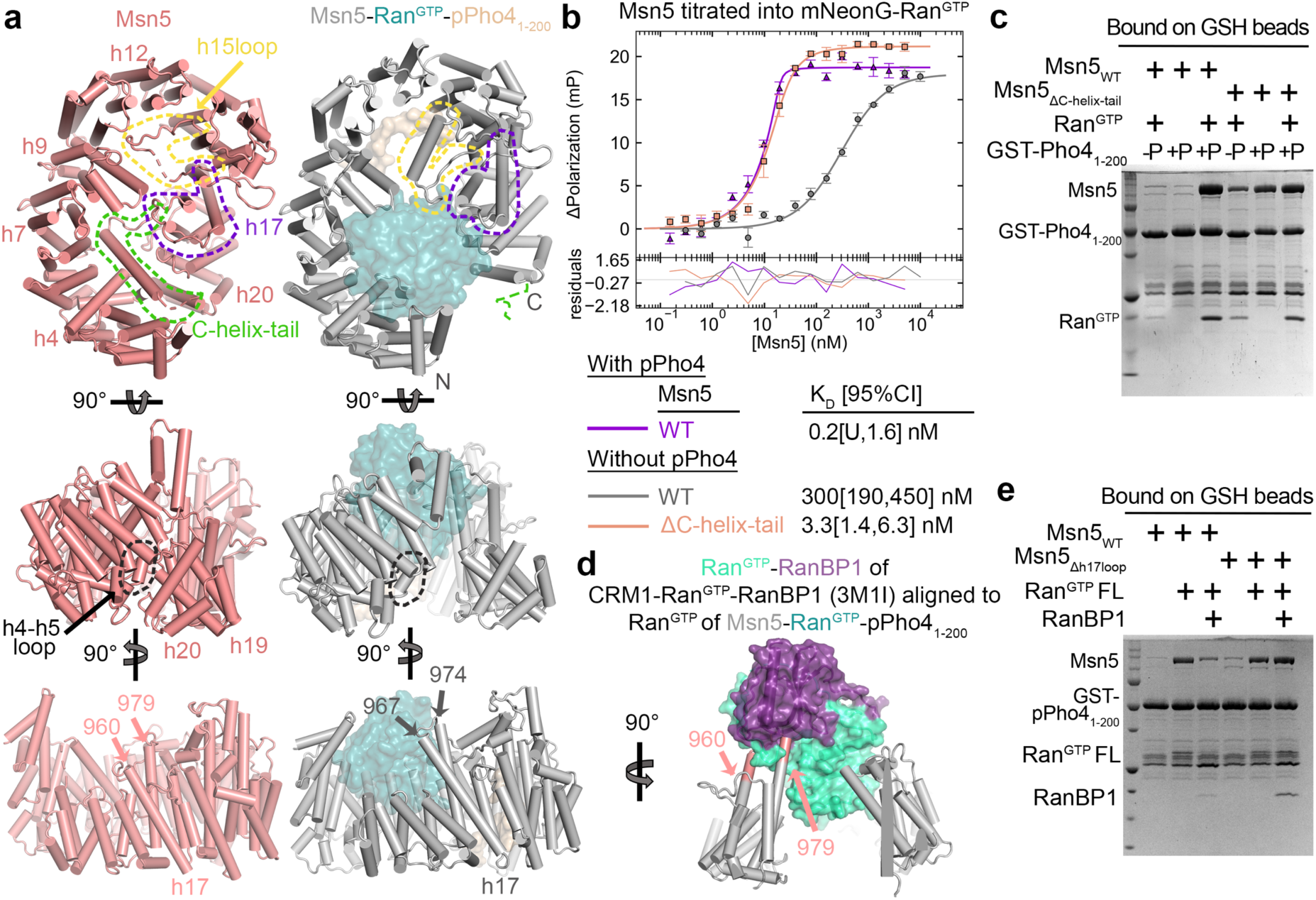
Unliganded Msn5 and the mechanisms for loading/unloading Pho4. **(a)** Unliganded Msn5 (pink, left) and Msn5-Ran^GTP^-pPho4_1-200_ (gray-teal-wheat, right). Structural differences at h15loop, h17 and C-helix-tail are highlighted by yellow, purple and green dashed lines, respectively. The h4-h5loop that contacts h19-h20 in unliganded Msn5 but not in ligand-bound Msn5, is highlighted with black dashed lines. (**b)** FP titration of WT or mutant Msn5_ΔC-helix-tail_ to mNeonG-Ran^GTP^ ± excess pPho4_1-200_. Data points are mean and s.d. of triplicate measurements, fitted with 1-site fitting (solid line) and fitting residuals plotted below. **(c)** Bound proteins (after extensive washing) in an *in vitro* pull-down assay of immobilized unphosphorylated (-P) or phosphorylated GST-Pho4_1-200_ (+P) incubated with Msn5 (WT or ΔC-helix-tail) ± Ran^GTP^, visualized by Coomassie stained SDS-PAGE. **(d)** Msn5 (gray; h17 residues 960-979 in pink)-Ran^GTP^-pPho4_1-200_ and CRM1-Ran^GTP^-RanBP1 (3M1I) structures are aligned at Ran^GTP^. CRM1 and pPho4 are not shown and the docked Ran-RanBP1 is aquamarine and purple, respectively. **(e)** As in **c**, pull-down assay of GST-pPho4_1-200_ incubated with Msn5 WT or Δh17loop (residues 962-980 replaced with GGSGGS) ± Ran^GTP^ FL ± RanBP1 as labeled. See also **Extended Data Fig. 10-12**.

Comparison of the Msn5-Ran^GTP^-pPho4_1-200_ and unliganded Msn5 structures shows that the Ran^GTP^-binding site at the N-terminal h1-h4 repeats of the former is occupied by HEAT repeats h17-h20 and the C-helix-tail in the latter (Fig. 6a, Extended Data Fig. 11a and b). Furthermore, the NES binding site (h8-h18) of unliganded Msn5 is substantially more compact, resulting in occlusion of the groove at h12-h15 that binds region 2 of the Pho4 NES (Extended Data Fig. 11b and c). These structural differences suggest that the unliganded conformation is autoinhibited and incompatible with ligand-binding. Interestingly, the dominant phospho-serine binding Msn5 RR site appears fully accessible, potentially allowing weak binding of part of the Pho4 NES (Extended Data Fig. 11c).

The autoinhibited conformation of unliganded Msn5 is consistent with weaker binding to either Ran^GTP^ or cargo alone. The K_D_ for Msn5 binding to mNeonG-Ran^GTP^ alone is 300 [190,450] nM compared to the sub-nanomolar K_D_ in the presence of pPho4_1-200_; Msn5 is also not pulled-down by immobilized GST-pPho4_1-200_ unless Ran^GTP^ is present (Fig. 6b and c, Extended Data Fig. 12a). These results show positive cooperativity of cargo and Ran^GTP^ binding, typical of exportin systems. The binding of either pPho4 or

Ran^GTP^ would stabilize Msn5 conformations with separated N/C-termini, increasing affinity for the other ligand (Extended Data Fig. 11a). The C-helix-tail that seems to hold together the termini of unliganded Msn5 may stabilize the autoinhibited state (Fig. 6a). Indeed, removal of the C-helix-tail (Msn5_ΔC-helix-tail_; residues 1179-1224 removed) increased Ran^GTP^ binding affinity by 100-fold (K_D_ 3.3[1.4, 6.3] nM for Msn5_ΔC-helix-tail_ *vs* K_D_ 300 nM for WT Msn5; Fig. 6b). Immobilized GST-pPho4_1-200_ also pulled-down Msn5_ΔC-helix-tail_ even when Ran^GTP^ is absent (Fig. 6c). Thus, the C-helix-tail of Msn5 is key for stabilizing its autoinhibited state.

Finally, we noticed that residues 960-979 of the h17loop of unliganded Msn5 adopt a different conformation in the Pho4/Ran^GTP^-bound Msn5. Here, residues 960-967 and 974-979 are helical and extend helices h17A and h17B, respectively, with the later segment contacting the Switch I of Ran^GTP^ (Fig. 6a and S4b). The Ran^GTP^ C-terminal tail is expected to become ordered upon binding the Ran^GTP^-binding protein RanBP1, which is known to be important for releasing cargoes from XPO1^51,52^. Alignment of Ran^GTP^-RanBP1 from the XPO1-bound structure (3M1I^51^) with the Ran^GTP^ of Msn5-Ran^GTP^-pPho4_1-200_ reveals a steric clash between the Msn5 h17loop-extended h17B helix with the superimposed Ran tail and RanBP1 (Fig. 6d). This observation suggests that RanBP1-binding might destabilize the Msn5-Ran^GTP^-pPho4 complex. Indeed, addition of RanBP1 (*Sc* Yrb1 Δ1-61) to Msn5 and Ran^GTP^ abolished binding to GST-pPho4_1-200_ (Fig. 6e). When the Msn5 h17loop residues, which are predicted to clash with Ran^GTP^-RanBP1, are replaced with a GGSGGS linker, the mutant Msn5_Δh17loop_ can be captured by GST-pPho4_1-200_ in complex with both Ran^GTP^ and RanBP1 (Fig. 6e). These results suggest that the h17loop of Msn5 is likely the allosteric switch that causes Ran^GTP^ release from Msn5 upon RanBP1-binding, and thus Pho4 (and other NES cargo) release.

To summarize, unliganded Msn5 adopts an autoinhibited conformation where its C-helix-tail closes the solenoid ring^50^. Ran^GTP^ binding displaces the C-helix-tail, opening the Msn5 solenoid to allow pPho4 to bind. Conformational changes of the Msn5 h17loop likely couples RanBP1-binding to Ran^GTP^ and cargo release to terminate a nuclear export cycle.

## Conclusion

We have identified a novel 35-residue phosphorylated Pho4 NES, the first member of a new class of NESs unrelated to the previously only known class of XPO1-recognized NESs, and explained how it is recognized by Msn5. The Pho4 NES has two critical phosphorylated serine residues and many small- to medium-sized hydrophobic and polar residues that all contribute to Msn5-binding. Knowledge of Msn5-binding elements and the recognition mode explains the mechanisms of nuclear export regulation by Pho4 phosphorylation. It also suggests criteria for other Msn5 cargoes even though the large and open Msn5 surface with multiple potential phospho-serine/threonine binding sites likely accommodates diverse NESs. The partially conserved binding surface of the homologous XPO5 may recognize even more divergent NESs. Finally, unliganded Msn5 is an autoinhibited closed-ring solenoid, and comparison with Ran^GTP^ and Pho4-bound Msn5 explains the positive cooperativity of Ran^GTP^ and cargo binding typical of exportins and cargo release by RanBP1.

## Acknowledgments

pL0M-S-mNeonGreen-EC18153 was a gift from Julian Hibberd (Addgene plasmid #137075; http://n2t.net/addgene:137075; RRID:Addgene_137075). pRSFDuet-1 vector was a gift from Christopher P. Hill. The ASY788 Msn5 knockout strain was a gift from Martha S. Cyert. We thank Yang Li, Yan Han and Zhe Chen at Structural Biology Laboratory, the CEMF in UTSW, and Sean Mulligan and Harry Scott at PNCC for their expert assistance with Cryo-EM data collection. We also thank the Erzberger lab for using their plate reader for FP. We thank Michael McConvile and Glen Liszczak for synthesizing phosphorylated peptides. SBL and CEMF at UTSW are partially supported by grant RP220582 from the Cancer Prevention & Research Institute of Texas (CPRIT). We thank the Genomics Resource Center at The Rockefeller University for assistance in mRNA sequencing library prep and data collection. A portion of this research was supported by NIH grant U24GM129547 and performed at the PNCC at OHSU and accessed through EMSL (grid.436923.9), a DOE Office of Science User Facility sponsored by the Office of Biological and Environmental Research. Molecular graphics and analyses performed with UCSF ChimeraX, developed by the Resource for Biocomputing, Visualization, and Informatics at the University of California, San Francisco, with support from National Institutes of Health R01-GM129325 and the Office of Cyber Infrastructure and Computational Biology, National Institute of Allergy and Infectious Diseases. This work was funded by NIGMS of NIH under Awards R35GM141461 (Y.M.C.), R01GM069909 (Y.M.C.), P41GM109824, R01 CA228351 and R01 GM112108 (M.P.R.), the Welch Foundation Grants I-1532 (Y.M.C.), support from the Alfred and Mabel Gilman Chair in Molecular Pharmacology, Eugene McDermott Scholar in Biomedical Research (Y.M.C.) and the Gilman Special Opportunities Award (H.Y.J.F.).

## Author contributions

Conceptualization, Y.M.C.; Methodology, Y.M.C., M.P.R., H.J.Y.F, S.R.M., A.E.C., T.Y.; Investigation, H.Y.J.F., S.R.M., A.N., J.J., B.S., A.E.C., T.Y.; Writing—Original Draft, H.Y.J.F., S.R.M.; Writing—Review & Editing, Y.M.C., H.Y.J.F, M.P.R., S.R.M.; Funding Acquisition, Y.M.C. and M.P.R.

## Methods

### Msn5 expression and purification

Msn5 was cloned into a pQE60 vector (C-terminal His) with the TEV cleavage sequence inserted and mutants and truncation mutants were generated by PCR and blunt-end ligation using WT vector. Msn5 was also cloned into pET28a vector with TEV cleavage site (N-terminal His). All Msn5 constructs were transformed into Rosetta 2(DE3) cells (Novagen) and grown in Luria Broth media at 37 °C with 100 mg/mL ampicilin and 25 mg/mL chloramphenicol until it reaches OD_600nm_ of 0.6. Expression was induced with 0.5 mM (for pQE60) or 0.25 mM (for pET28a) IPTG for 12 hr at 18 °C. Cells were spun down and resuspended with buffer A containing 50 mM HEPES pH 7.4, 500 mM sodium chloride, 10 % (v/v) glycerol, supplemented with 2 mM 2-mercaptoethanol, 1 mM benzamidine, 10 μg/mL leupeptin and 50 μg/mL AEBSF before use. The resuspension was lyzed with an EmulsiFlex-C5 (Avestin) at ∼ 10,000 psi and clarified by centrifugation at 20,000 rpm. The supernatant was supplemented with 10 mM imidazole pH 7.8 and incubated with nickel beads (Qiagen) for 10 – 30 min. Beads were washed with buffer A supplemented with 10 mM imidazole, 2 mM 2-mercaptoethanol and Msn5-His was eluted with buffer A containing 500 mM imidazole, 2 mM 2-mercaptoethanol. The elution was loaded onto HiTrap Phenyl HP column (Cytiva) and eluted with buffer containing 50 mM HEPES pH 7.4, 1 mM DTT and 10 % glycerol. 1 mg of recombinant purified Tev was added to Msn5-His and incubated overnight at 4 °C. Tev-cleaved Msn5 was further purified in Superdex 200 Increase 10/300 GL column (Cytiva) in buffer B containing 20 mM HEPES pH 7.4, 150 mM sodium chloride, 10 % glycerol and 2 mM TCEP, and flash-frozen in liquid nitrogen and stored in −80 °C.

### Ran^GTP^ constructs and purification

Ran^GTP^ is expressed using pET21d-Ran^GTP^ construct containing yeast Ran, Gsp1, residues 1-179 with Q71L mutation to stabilize the GTP bound form, and no manual GTP loading is needed. It is expressed and purified as described previously with the addition of TEV cleavage.^53^ Briefly, Ran^GTP^-His was expressed in BL21(DE3) cells (New England Biolabs) using 0.5 mM IPTG for 12 hr at 20 °C. Cells were resuspended in buffer containing 50 mM HEPES pH 7.4, 2 mM magnesium acetate, 200 mM sodium chloride, 10 % glycerol, 5 mM imidazole pH 7.8, 2 mM 2-mercaptoethanol, 1 mM benzamidine, 10 μg/mL leupeptin, 50 μg/mL AEBSF. It was then lyzed and clarified by centrifugation, and the supernatant was incubated with nickel beads before it was washed and eluted with 50 mM HEPES pH 7.4, 2 mM magnesium acetate, 50 mM sodium chloride, 10 % glycerol, 250 mM imidazole pH 7.8, 2 mM 2-mercaptoethanol. The elution was concentrated, and 1 mg of TEV was added for incubation overnight at 4 °C. TEV-cleaved Ran^GTP^ was purified by ion-exchange using HiTrap SP HP column (Cytiva) over a gradient of 25 to 500 mM sodium chloride and clean protein was flash-frozen and stored in −80 °C.

Ran^GTP^ and mNeonGreen (from pL0M-S-mNeonGreen-EC18153) were inserted into pET28a to generate pET28a-tev-mNeonG-Ran^GTP^ construct. His-mNeonG-Ran^GTP^ was purified using Rosetta(DE3)pLysS cells (Novagen) grown with 50 mg/mL kanamycin and 25 mg/mL chloramphenicol. Expression was induced with 0.25 mM IPTG at 18 °C for 12 hr. Purification was performed like TEV-cleaved Ran^GTP^ to cleave off the His tag.

Ran^GTP^ FL-His was expressed using pET15b-Ran^GTP^ FL in BL21 Gold(DE3) cells with 0.5 mM IPTG for 3.5 hr at 37 °C and purified as in human Ran FL in an established protocol.^54^ Purification is similar to Ran^GTP^ but the His tag is left intact and protein is subjected to final purification in Superdex 75 10/300 column (Cytiva) in Buffer B instead. GTP is loaded on Ran^GTP^ FL by incubating the protein with 2 mM GTP and 10 mM EDTA pH 8.0 for 40 min on ice, and addition of 40 mM magnesium acetate directly before use.

### Purification of RanBP1

GST-RanBP1 was expressed in BL21(DE3) cells using pGex-tev-RanBP1 (yeast Yrb1, residues 61-201) with 0.5 mM IPTG at 25 °C for 10 hr and purified according to established protocol.^53^ In short, cells were lyzed in buffer containing 40 mM HEPES pH 7.5, 2 mM magnesium acetate, 200 mM sodium chloride, 10 mM DTT, 1 mM benzamidine, 10 μg/mL leupeptin, 50 μg/mL AEBSF and clarified, and supernatant was bound to glutathione Sepharose beads and washed with 100 mM and 300 mM sodium chloride. GST-RanBP1 was eluted with 30 mM GSH and cleaved with TEV overnight at 4 °C. Reaction was passed over GSH beads again before it was purified over Superdex 75.

### Pho4 constructs and purification

Full length and residues 1-200 of Pho4 were cloned from yeast cDNA library into pGex-tev construct. mNeonGreen (mNeonG) and Pho4(3-200) were both inserted into pET28a vector to generate pET28a-tev-mNeonG-Pho4_3-200_ construct. All mutants were generated by site-directed mutagenesis. All GST constructs were transformed into BL21(DE3) cells and grown in LB with 100 mg/mL ampicillin. Expression was induced with 0.5 mM IPTG at 25 °C for 12 hr for GST-Pho4_FL_ or 30 °C for 3.5 hr for GST-Pho4_1-200_ variants. mNeonG-Pho4 variants were expressed like mNeonG-Ran^GTP^.

Cells with GST proteins were lysed in buffer A supplemented with 2 mM DTT, 1 mM benzamidine, 10 μg/mL leupeptin and 50 μg/mL AEBSF, clarified and bound to glutathione Sepharose beads (Cytiva), with the exception of GST-Pho4_FL_ where HEPES pH 7.5 was replaced with Tris pH 8.0. Beads were washed with the same buffer and eluted with buffer containing 25 mM sodium chloride and supplemented with 30 mM GSH. GST-Pho4_FL_ was further purified by Hi Trap Q HP column (Cytiva) in 20 mM Tris pH 8.0, 10 % glycerol, 2 mM DTT using a gradient of 25 mM – 1 M sodium chloride. To generate Pho4^FL^, beads were washed with buffer containing 20 mM Tris pH 8.0, 10 % glycerol, 150 mM sodium chloride, 1 mM DTT and 1 mg TEV was added to perform cleavage on-beads for 2 hr at room temp. Flowthrough was purified in Q column like GST-Pho4_FL_ but in Tris pH 8.5. GST-Pho4_1-200_ eluted from beads was purified by Q in HEPES 7.0. Pho4_1-200_ is generated in a similar manner as Pho4_FL_ but the Q is performed in HEPES pH 7.4.

Cells expressing His-mNeonG-Pho4_1-200_ proteins were lyzed in buffer A in Tris pH 8.0 with 10 mM imidazole pH 7.8, 2 mM 2-mercaptoethanol, 1 mM benzamidine, 10 μg/mL leupeptin and 50 μg/mL AEBSF and clarified. Supernatants were purified over HisTrap HP (Cytiva) in 50 mM Tris pH 8.0, 150 mM sodium chloride, 10 % glycerol, 10 – 150 mM imidazole pH 7.8, 2 mM 2-mercaptoethanol. Fractions containing His-mNeonG-Pho4_1-200_ were concentrated and cut with Tev overnight to remove the His tag at 4 °C before further purification over Q column in HEPES pH 7.0 like the GST proteins and frozen.

### In vitro phosphorylation of Pho4

His-Pho85 and Pho80 were cloned into MCS-1 and 2 respectively in pRSFDuet vector and was expressed in Rosetta(DE3)pLys cells using 1 mM IPTG at 18 or 25 °C for 12 hr. Cells were lysed in buffer containing 20 mM HEPES pH 7.4, 150 mM sodium chloride, 10 % glycerol, 1 mM 2-mercaptoethanol and 40 mM imidazole pH 7.8, then clarified, and the supernatant was incubated with nickel beads. Beads were washed with 80 mM Imidazole and His-Pho85/Pho80 complex was eluted with 300 mM Imidazole. The complex was further purified over Superdex Increase S200 in buffer containing 20 mM HEPES pH 7.4, 150 mM sodium chloride, 10 % glycerol, 1 mM DTT and 10 mM magnesium chloride.

10 – 30 µM of Pho4 variants were mixed with 0.5 mM His-Pho85/Pho80 and ATP at 300 times the substrate concentration in 50 mM HEPES pH 7.4, 150 mM sodium chloride, 1 mM DTT, 10 mM magnesium chloride, 10% glycerol, and incubated at room temp for 1 hour. Proteins for pulldown binding assays were bound to beads and washed, while proteins for fluorescence polarization measurements were buffer exchanged on a Superdex 75 into buffer B. Samples before and after phosphorylation was sent to intact mass analysis to ensure that phosphorylation was complete.

### In vitro pull-down binding assays

For Fig. 1, 1 µM Msn5, 5 µM Ran^GTP^ and ∼1-2 µM GST proteins were mixed in buffer B in 200 µL reactions with ∼10-20 µL glutathione Sepharose beads and rotated at 4 °C for 30 mins. Samples were then spun at 6000 xg for 1 min and flowthrough was removed. Buffer B was used to wash the beads for 3 times, 500 µL each. Lastly, 20 µL 2X SDS sample buffer was added to beads and boiled for 3 mins and bound proteins were visualized on SDS-PAGE gel for Coomassie staining. ∼2.5 µM GST-Pho4_1-200_ or GST-pPho4_1-200_ + 1 µM Msn5 (WT or mutants) ± 5 µM Ran^GTP^ for assay in Fig. 6c, or ∼2.5 µM GST-pPho4_1-200_ + 1 µM Msn5 ± 5 µM Ran^GTP^ FL ± 5 µM RanBP1 for assay in Fig. 6e, were assembled and treated the same way as described above. For binding assay in Extended Data Fig.1a, GST-Pho4 proteins were expressed and purified in small scale (35 mL). Lysate was sonicated over nice before they were clarified and bound to GSH beads and washed. Beads were used directly for binding assay with 5 µM Msn5 and 20 µM Ran^GTP^ following procedures described above.

### Cryo-EM sample preparation and data collection

Complexes were formed by mixing Msn5, Ran^GTP^ and phosphorylated Pho4 in 1: 10:2 ratio and purified over Superdex S200 Increase in 20 mM HEPES pH 7.4, 150 mM sodium chloride, 2 mM magnesium acetate, 2 mM TCEP. Msn5-Ran^GTP^-pPho4_1-200_ was diluted to 3.7 mg/mL and supplemented with 0.05 % NP-40 (Thermo-Fisher). Msn5-Ran^GTP^-pPho4_FL_ was diluted to 3 mg/mL and supplemented with 0.00125 % NP-40. Msn5 was diluted to 6 mg/mL in 20 mM HEPES pH 7.4, 150 mM sodium chloride, 2mM TCEP with a final concentration of 0.1 % NP-40. Samples were applied on holey carbon grids (Quantifoil R1.2/1.3, 300 mesh copper) that were previously glow-discharged using a PELCO easiGlow glow discharge apparatus for 60 s at 30 mA (Ted Pella), and then plunge frozen using the Vitrobot Mark IV System (Thermo Fisher). One grid with best particle distribution for pPho4_1-200_-containing complex was used for 36 – 48 hr data collection on a Titan Krios microscope equipped with K3 detector at the Cryo Electron Microscopy Facility (CEMF) at UT Southwestern Medical Center. Two different grids for pPho4_FL_-containing complex were shipped to Pacific Northwest Cryo-EM Center (PNCC) for 72 hr data collection on the Titan Krios microscope equipped with K3 detector. A total of 13,496 movies were acquired from one grid at pixel size of 0.5115 Å in super-resolution counting mode. Two different Msn5 grids with best particle distribution were shipped to PNCC for 48 h data collection. A total of 8490 movies were acquired from the two grids collected the same way as the pPho4_FL_ sample.

### Cryo-EM data processing

All data processing was performed using the software cryoSPARC v3.3.1 and v3.3.2 using default parameters^55^. A ½ F-crop factor was applied during patch motion correction, followed by patch CTF estimation. For Msn5-Ran^GTP^-pPho4_1-200_, blob picker was used to select 3,752,564 initial particles, which cleaned up to 1,574,329 particles after one round of 2D classification. After additional 9 extensive rounds of 2D classification, 883,743 particles were used to obtain 3 *ab initio* models, which were subsequently submitted to heterogeneous refinement. ∼30% of the particles partitioned into a map that looks like Msn5 alone. The rest of the particles, a total of 617,333 particles can be further refined using non-uniform refinement to generate a 2.97 Å resolution map which showed clearly density for Ran^GTP^ but not Pho4. These particles were then further used to generate 6 new *ab initio* models which were then submitted to heterogenous refinement and non-uniform refinement with per-particle defocus and per-group CTF optimization.

Msn5-Ran^GTP^-pPho4_FL_ was processed similarly and had 1,104,150 particles after a single round of 2D classification. After 7 additional rounds of 2D classification, 395,996 particles were classified into 6 classes, only 2 of which with most particles looked like good particles and were submitted to non-uniform refinement without per-particle defocus and per-group CTF optimization to yield two maps of ∼ 5 Å resolution.

For unliganded Msn5, blob picker was used to select initial particles which cleaned up to 4,135,768 after one 2D classification run. After 6 rounds of 2D classification, 1,450,905 particles were used to generate 6 *ab initio* models. Some junk classes were observed, therefore more particles (1,998,715 particles from one less round of 2D classification) were submitted for heterogenous refinement with three input models which included one class to sink junk particles. The best map of 3.39 Å resolution was obtained by further non-uniform refinement of a subset of 1,048,421 particles, with per-particle defocus and per-group CTF refinement. Including the other subset of particles (726,480 particles), which reconstructed into a highly similar map, did not improve resolution.

### Cryo-EM model building, refinement and analysis

Unsharpened map was used for model building as they have the best overall features. SWISS-MODEL was used to generate an initial 3D-model of Msn5 using Exportin-5 structure as a template (PDB ID: 5YU7). The Msn5 model and Ran^GTP^ from PDB ID 3M1I were fitted into the cryo-EM map using UCSF ChimeraX^56^, followed by multiple cycles of manual model building using ISOLDE^57^ in ChimeraX, Coot^58^ and real-space refinement in Phenix^59^. Maps often have poor density for the last two HEAT repeats at the C-terminal, and the models are all generated by merging with a fragment of a docked model of Msn5 alone. Model validation was performed in Phenix. The binding interfaces were analyzed using CONTACT/ACT in CCP4 with cut-off of 4.0 Å^60^. These interactions were curated and analyzed in PyMOL 2.4.2 where the final figures were generated^61^. Map images were generated in ChimeraX.

### Fluorescence Polarization assays

Fluorescence polarization assays were performed in the same way as previously described.^53^ In brief, triplicates of sixteen 20 μL samples were assembled with final concentration of 20 nM mNeonG-pPho4_1-200_, 30 μM Ran^GTP^ and a 1:1 serially diluted wild type or mutant Msn5 from 10 μM, in a 384 well black bottom plate (Corning). For Ran^GTP^ binding affinities, samples contain 20 nM mNeonG-Ran^GTP^ with serial dilution of wild type or mutant Msn5 from 10 μM. Measurements were performed in a CLARIOstar Plus plate reader (BMG Labtech) with top optics using excitation filter 482-16, dichroic filter LP 504 and emission filter 530-40, 50 flashes per well. Gain was optimized to target mP of ∼220 for mNeonG-pPho4_1-200_ and ∼200 for mNeonG-Ran^GTP^. Data was analyzed in PALMIST^62^ and plotted in GUSSI^63^.

### RNAseq sample preparation, data collection and processing

Msn5 was cloned into a p413 yeast expression vector (His selection) with a Cyc1 promoter inserted and the HRYRR_AAAAA_ mutants were generated by PCR using WT vector. The ASY788 strain (Msn5 knockout) was acquired from Martha S. Cyert’s Lab and transformed with either an empty p413 vector, a p413 WT Msn5 vector, or a p413 HRYRR_AAAAA_ Msn5 mutant vector. Colonies were assessed for proper incorporation by plating on synthetic complete medium agar plates without Histidine, so only cells carrying the plasmids conferring the His selection marker would grow. Single colonies were taken directly off the plate and inoculated into 5mL overnight cultures and grown to mid-log phase. Cultures were spun down, washed twice with PBS, pelleted and flash frozen prior to downstream processing. Each strain was grown in triplicate.

Total RNA from each pellet was extracted using the Monarch Total RNA Miniprep Kit. In brief, pellets were resuspended in 1X DNA/RNA protection reagent, glass beads were added to the resuspension before vortexing at high speed to lyse. Samples were spun down and supernatant was spun through a column to remove gDNA. Flowthrough was mixed with ethanol to facilitate binding to an RNA purification column, onto which samples were subjected to DNase I treatment, before being washed several times and eluted in nuclease-free water. Illumina mRNA-seq library preps were conducted by The Rockefeller University Genomics core facility. Samples were pooled and analyzed with a NextSeq 2000 instrument with high output flowcell 75 single read sequencing.

Initial sequencing data was processed in Linux, starting with the FastQC software used to check the quality of the raw data. Poor quality regions and adapter sequences were trimmed using Trimmomatic. Reads were aligned to the *Saccharomyces cerevisiae* genome (acquired from *Saccharomyces* Genome Database) using STAR. Finally gene hit counts were determined using FeatureCounts. Comparison of hit counts between conditions and any further downstream analysis was conducted using R scripting. Comparison between conditions was conducted using DESeq2 and volcano plots were generated using ggplot2.

