## Supplemental Information for "Phosphate-dependent nuclear export via a novel NES class recognized by exportin Msn5"

**Analysis of Pho4 density/map features in the additional cryo-EM maps of the Msn5-$Ran^{GTP}$-$pPho4_{1-200}$ complex**

We divided Msn5-$Ran^{GTP}$-$pPho4_{1-200}$ particles into 3 groups named State 1, State 2 and State 3 according to the different extent of compact to open curvatures of their Msn5 solenoids. The Msn5 in the State 1 is most compact and most open in State 3. Each state contains two populations (e.g., State 1-1 and State 1-2) with small differences in Msn5 solenoid conformations (Extended Data Fig. 2a). Cryo-EM maps features for Pho4 NES can be observed to varying degrees in all six cryo-EM maps generated from the six group of particles (Extended Data Fig. 4b-d). We focused on describing the most compact State 1-1 map of Msn5-$Ran^{GTP}$-$pPho4_{1-200}$ and its resulting structural model in the main text. Here, we analyze the differences in the additional maps that show the multiple conformations and structural flexibility of Msn5.

In the State 1-1 map of Msn5-$Ran^{GTP}$-$pPho4_{1-200}$, which has the most compact Msn5 conformation, density for the Pho4 NES across regions 2 to 4 is continuous and well-defined. In the slightly less compact State 1-2 map, density for the pPho4 NES region 2 is also continuous but not as well-defined as in the State 1-1 map (Extended Data Fig. 4b). In the States 2-1, 2-2, 3-1 and 3-2 maps of Msn5-$Ran^{GTP}$-$pPho4_{1-200}$, Pho4 densities are no longer continuous. The best defined Pho4 features of these maps lie adjacent to Msn5 repeats h8-h10, including the Msn5 RR site, where Pho4 NES region 4 ($^{128}$pSPNL$^{131}$) binding can be observed in the States 1-1 and 1-2 maps (Extended Data Fig. 4c and d). However, the features at the same location in the State 2-1, 2-2, 3-1 and 3-2 maps seem to fit better to $^{113}$ApSPLA$^{117}$ than to $^{128}$pSPNL$^{131}$ (Extended Data Fig. 13a). The map features are too ambiguous to confidently model Pho4 but raise the possibility that the Msn5 RR site may also engage pS114. Nonetheless, the maps suggest that a phosphoserine containing segment of the Pho4 NES likely always occupies the Msn5 RR site, consistent with mutagenesis data suggesting that Msn5 RR is the more important phosphoserine binding site (Fig. 3b).

Several maps also contain extra density sandwiched between $Ran^{GTP}$ and the h6-h8 repeats of Msn5 (Extended Data Fig. 4). Mutation of Msn5 residues here, D285, H288, D343 to alanines, did not affect binding to $Ran^{GTP}$ or to $pPho4_{3-200}$ by FP (Extended Data Fig. 13b). It is possible that this density is the NP-40 detergent.

**Analysis of Pho4 map features in the additional cryo-EM map of the Msn5-$Ran^{GTP}$-$pPho4_{FL}$ complex**

Like the Msn5-$Ran^{GTP}$-$pPho4_{1-200}$ particles, the Msn5- $Ran^{GTP}$-$pPho4_{FL}$ particles also displayed flexibility. Two reconstructions were obtained, which matched the Msn5 solenoid shape of State 1 and 3 maps from the $pPho4_{1-200}$ dataset (Extended Data Fig. 2a). The map features for Pho4 NES is continuous in the State 1 map (most compact Msn5) of $pPho4_{FL}$-bound Msn5 but not continuous for the State 3 map (most open Msn5 conformation; Extended Data Fig. 4e). The State 3 map has map features for Pho4 NES regions 1 and 4 but not for regions 2 and 3. Interestingly, NES region 1 of $pPho4_{FL}$ appears persistently bound to Msn5 even though the same region of $pPho4_{1-200}$ is not observed bound to Msn5 (Extended Data Fig. 4). It is possible that additional $pPho4_{FL}$ residues that are not present in the shorter $pPho4_{1-200}$ construct may engage Msn5, influencing the more persistent interaction of NES region 1 of $pPho4_{FL}$ with Msn5. Alternatively, these additional $pPho4_{FL}$ residues include a putative oligomerization domain at residues 201-300, which may facilitate Pho4 dimerization/oligomerization, influencing Pho4 NES accessibility and Msn5 binding. The 2-site binding curve of mNeonG-pPho4$_{FL}$ for Msn5 suggests that when pPho4 dimerizes, one pPho4 subunit is less accessible to Msn5 when the other subunit is already bound to Msn5 (Extended Data Fig. 1b).

**Comparison of pPho4 S114A and S128A mutants and the Msn5 mutants at their respective binding sites**

As described in the main text, we mutated the Pho4 serines that are phosphorylated (Pho4 S114$_A$, Pho4$_{1\text{-}200}$ S128$_A$ and Pho4$_{1\text{-}200}$ S114/128$_{AA}$) and the phosphate binding sites on Msn5 (Msn5 HRY$_{AAA}$ and Msn5 RR$_{AA}$). The 5-fold affinity decrease of the Msn5 HRY$_{AAA}$ mutant ($K_D$ 230[160,350 nM) compared to Msn5 WT matches the 6-fold decrease seen for pPho4 S114$_A$ vs. pPho4 WT. However, the 50-fold affinity decrease of Msn5 RR$_{AA}$ ($K_D$ 2.2[1.0,6.1] μM) vs the 3-fold decrease of pPho4 S128$_A$ is surprising as Msn5 R393 and R458 contact only the phosphate group of pS128 in the structure (Fig. 2c, left panel). It is possible that the loss of the phosphate group in pPho4 S128$_A$ is less detrimental because Msn5 R393 and R458 may still interact with the alanine at position 128 or its neighboring residues. Alternatively, in the absence of pS128, pS114 may bind at the Msn5 RR site (see above).

**Interactions of pPho4 S114A and S128A with Msn5 HRY$_{AAA}$ and RR$_{AA}$ mutants**

Next, we combined Pho4 and Msn5 mutations to validate the pairing of Pho4 pS114 binding at Msn5 HRY and pS128 at Msn5 RR. Consistent with the former pair, the affinity loss of pPho4 binding to Msn5 HRY$_{AAA}$ (vs WT) is not exacerbated by additionally mutating S114 ($K_D$ to Msn5 HRY$_{AAA}$ = 300[170,520] nM for S114$_A$ *vs.* 230[160,350] nM for WT; Extended Data Fig. 6d). The same is observed for the latter pair ($K_D$ to Msn5

$RR_{AA}$ = 2.8[2.4,3.1] μM for $S128_A$ *vs.* 2.2[1.0,6.1] μM for WT; Extended Data Fig. 6e). In contrast, mutations involving mispaired sites, pPho4 $S114_A$ binding Msn5 $RR_{AA}$ or pPho4 S128A binding Msn5 $HRY_{AAA}$, showed additional affinity decreases (Extended Data Fig. 6d and e). In summary, mutagenic analysis is consistent with the cryo-EM structure which shows Pho4 pS114 binding at the Msn5 HRY site, and pS128 binding at the Msn5 RR site.

**Extended Data Table 1. Cryo-EM data and map statistics of additional maps from pPho4 complexes.**

| Complex | Msn5-Ran$^{GTP}$-pPho4$_{1-200}$ | | | | |
|---|---|---|---|---|---|
| **State** | **1-2** | **2-1** | **2-2** | **3-1** | **3-2** |
| Magnification | 105,000 | | | | |
| Voltage | 300 kV | | | | |
| Electron exposure | 50 e$^{-}$/Å$^{2}$ | | | | |
| Defocus range | -0.8 to -2.5 μm | | | | |
| Pixel size | 0.83 Å | | | | |
| Symmetry imposed | C1 | | | | |
| Initial particle images[1] | 1,574,329 | | | | |
| Final particle images | 93,291 | 102,289 | 85,048 | 112,923 | 121,812 |
| Map resolution (Å)[2] | 3.37 | 3.19 | 3.33 | 3.11 | 3.23 |
| *B* factor (Å$^{2}$) | 92.2 | 90.7 | 81.8 | 89.3 | 100.2 |
| EMDB ID | EMD-46557 | EMD-46558 | EMD-46560 | EMD-46551 | EMD-46556 |

| Complex | Msn5-Ran$^{GTP}$-pPho4$_{FL}$ | |
|---|---|---|
| **State** | **1** | **3** |
| Magnification | 81,000 | |
| Voltage | 300 kV | |
| Electron exposure | 50 e$^{-}$/Å$^{2}$ | |
| Defocus range | -0.8 to -2.5 μm | |
| Pixel size | 1.03 Å | |
| Symmetry imposed | C1 | |
| Initial particle images[1] | 1,104,150 | |
| Final particle images | 108,248 | 111,752 |
| Map resolution (Å)[2] | 4.91 | 5.12 |
| *B* factor (Å$^{2}$) | 325.9 | 387.4 |
| EMDB ID | EMD-46562 | EMD-46561 |

[1]Number after 1$^{st}$ round 2D to remove images that are clearly not protein particles.
[2]FSC threshold = 0.143

**Extended Data Table 2. Segments of other Msn5 cargoes with similar characteristics as the Pho4 NES.** These characteristics include phospho-sites >10 residues apart, the presence of many hydrophobic and small polar side chains and few basic side chains.

| Cargo | Putative NES region | Sequence* |
|---|---|---|
| Crz1 | 186-279 (sufficient to direct export)[1] | 186-SSGID SNYSDTESNY HTPYLYPQDL VSSPAMSHLT ANNDDFDDLL SVASMNSNYL PVNSHGYKH<br>251 ISNLDELDDL LSLTYSDNNL LSASNNSDF |
| Swi5 | 1-325 (smallest fragment interacting with Msn5 in yeast two-hybrid analysis (Y2H))[2] | 1 MDTSNSWFDA SKVQSLNFDL QTNSYYSNAR GSDPSSYAIE GEYKTLATDD LGNILNLNYG ETNEVIMNEI<br>71 NDLNLPLGPL SDEKSVKVST FSELIGNDWQ SMNFDLENNS REVTLNATSL LNENRLNQDS GMTVYQKTMS<br>141 DKPHDEKKIS MADNLLSTIN KSEINKGFDR NLGELLLQQQ QELREQLRAQ QEANKKLELE LKQTQYKQQQ<br>211 LQATLENSDG PQFLSPKRKI SPASENVEDV YANSLSPMIS PPMSNTSFTG SPSRRNNRQK YCLQRKNSSG<br>281 TVGPLCFQEL NEGFNDSLIS PKKIRSNPNE NLSSKTKFIT PFTPK |
| Aft1[†#] | 147-270, 304-498 (either segment interacted with Msn5 in Y2H, but both were needed for export)[3] | 147-KRGR NARRKRKDKP KGQDHEDEKS KINDDELEYA SPSNATVTNG PQTSPDQTSS IKPKKKRCVS<br>211 RFNNCPFRVR ATYSLKRKRW SIVVMDNNHS HQLKFNPDSE EYKKFKEKLR KDNDVDAIKK<br>304-VVLPTNS NVTSSASSST VSSISLDSSN ASKRPCLPSV NNTGSINTNN VRKPKSQCKN KDTLLKRTTM<br>371 QNFLTTKSRL RKTGTPTSSQ HSSTAFSGYI DDPFNLNEIL PLPASDFKLN TVTNLNEIDF TNIFTKSPHP<br>441 HSGSTHPRQV FDQLDDCSSI LFSPLTTNTN NEFEGESDDF VHSPYLNSEA DFSQILSS |
| Mig1[†] | 244-340 (sufficient to direct export)[4] | 244-RTVFIDG PEQKQLQQQQ NSLSPRYSNT VILPRPRSLT DFQGLNNANP NNNGSLRAQT QSSVQLKRPS<br>311 SVLSLNDLLV GQRNTNESDS DFTTGGEDEE |
| Rtg3 | 2-279 (removal caused nuclear retention)[5] | 1 MMNNNESEAE NQRLLDELMN QTKVLQETLD FSLVTPTPHH NDDYKIHGSA YPGGETPAQQ HEKLSYINTH<br>71 NSNDNNNLMG SQARSNSQTP TASTIYEEAE SQSSYLDDMF RTSQGGRPVT QNSISSIGQG PLRSSYSMAY<br>141 DSPVDRAMNT PLQQQEGLKA ELPHDFLFQH GTDDTMYNLT DDLSSSLSSS INSDMMTPNT YSSSFSYNPQ<br>211 SLGPASVSST YSPKVRSPSS SFRAGSFLSS SFRHGSINTP RTRHTSISSN MTENIGPGSV PKILGGLTS |
| Cdh1 | 1-50 (deletion mutant did not interact with Msn5 in Y2H)[6] | 1 MSTNLNPFMN NTPSSSPLKG SESKRVSKRP ISSSSSASLL SSPSRRSRPS |
| Far1 | 285-390 (deletion mutant did not interact with Msn5 in Y2H and caused nuclear retention)[7] | 285-MQQQWI DLKTARSFTG EFPQFTPQEQ LIRTADISCD GFRTPRLSNS NQFEAVSYLD SPFLNSPFVN<br>351 KMATTDPFDL SDDEKLDCDD EIDESAAEVW FSKTGGEHVM |
| Ste5[§] | 24-55, 160-214 (deletion mutants did not interact with Msn5 in Y2H and caused nuclear retention)[8] | 24-GTLSRTP NQIIELEKPS TLSPLSRGKK WTEKL<br>160-G YPIQRTSIKK SFLNASCTLC DEPISNRRKG EKIIELACGH LSHQECLIIS FGTTSKAD |

* In bold, residues marked by SGD as phosphorylation sites.

[†] In red, mutations of S210/S224, T421/T423/T431/435 or S210/S224/T421/T423/T431/435 to alanines abolished export.

[#] Aft1 S210A/S124A mutant did not interact with Msn5 (Y2H). Aft1 T421A/T423A/T431A/435A mutant interacted with Msn5 (Y2H).

[§] In green, Ste5 T29A/S43A (together with other mutations outside this region) reduced Msn5 interaction (Y2H).

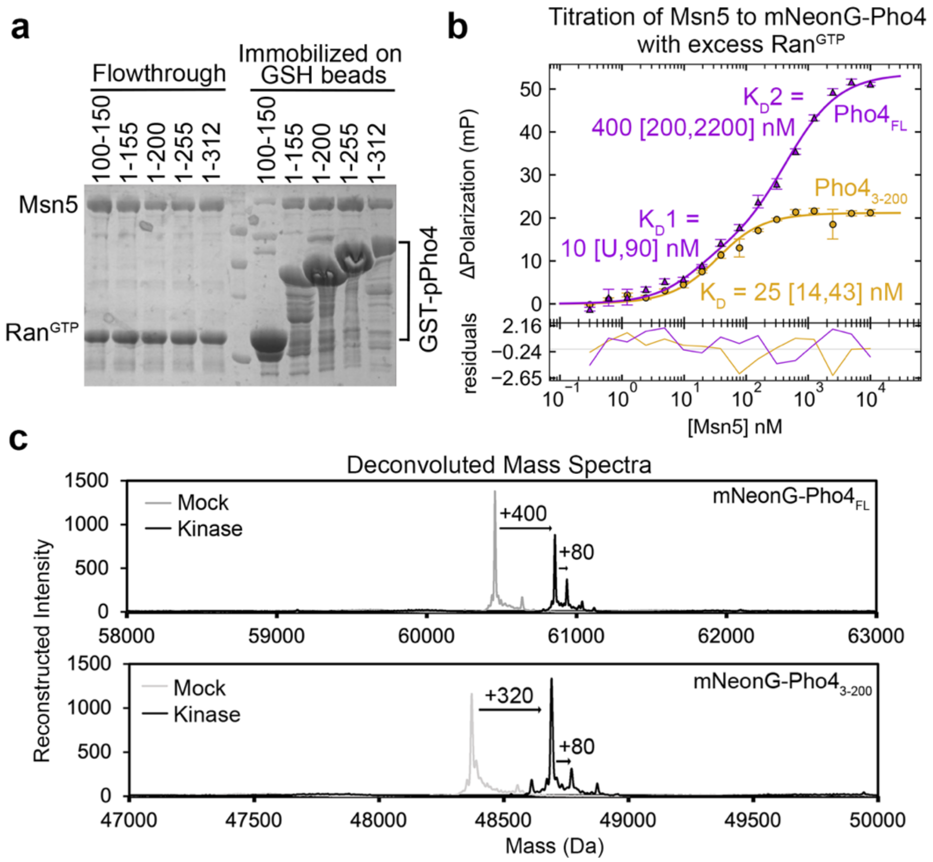


**Extended Data Fig. 1. Biochemical analysis of Pho4 binding to Msn5. (a)** GST-Pho4 constructs of different lengths were phosphorylated and then immobilized on GSH beads, incubated with Msn5 and $Ran^{GTP}$ and washed extensively before the unbound (flowthrough) and bounds proteins were visualized by SDS-PAGE and Coomassie staining. **(b)** FP titration of Msn5 into 20 nM mNeonG-Pho4 (FL or 3-200) in the presence of 30 μM $Ran^{GTP}$**.** Data points are averages of triplicate measurements with error bars representing standard deviation. $Pho4_{FL}$ data is fitted using two-site (independent) binding model and assuming mNeonG-$Pho4_{FL}$ is at 10 nM dimer concentration, whereas $Pho4_{3\text{-}200}$ data is fitted with 1-site binding model. Fitting residues are plotted below. Dissociation constants ($K_D$) are reported with 95% confidence interval in brackets. U means unable to be determined. **(c)** Deconvoluted mass spectra from intact mass analysis of mNeonG-Pho4 proteins used in **(b)**. mNeonG-$Pho4_{FL}$ contains at least 5 phosphates and mNeonG-$Pho4_{3\text{-}200}$ has 4 phosphates.

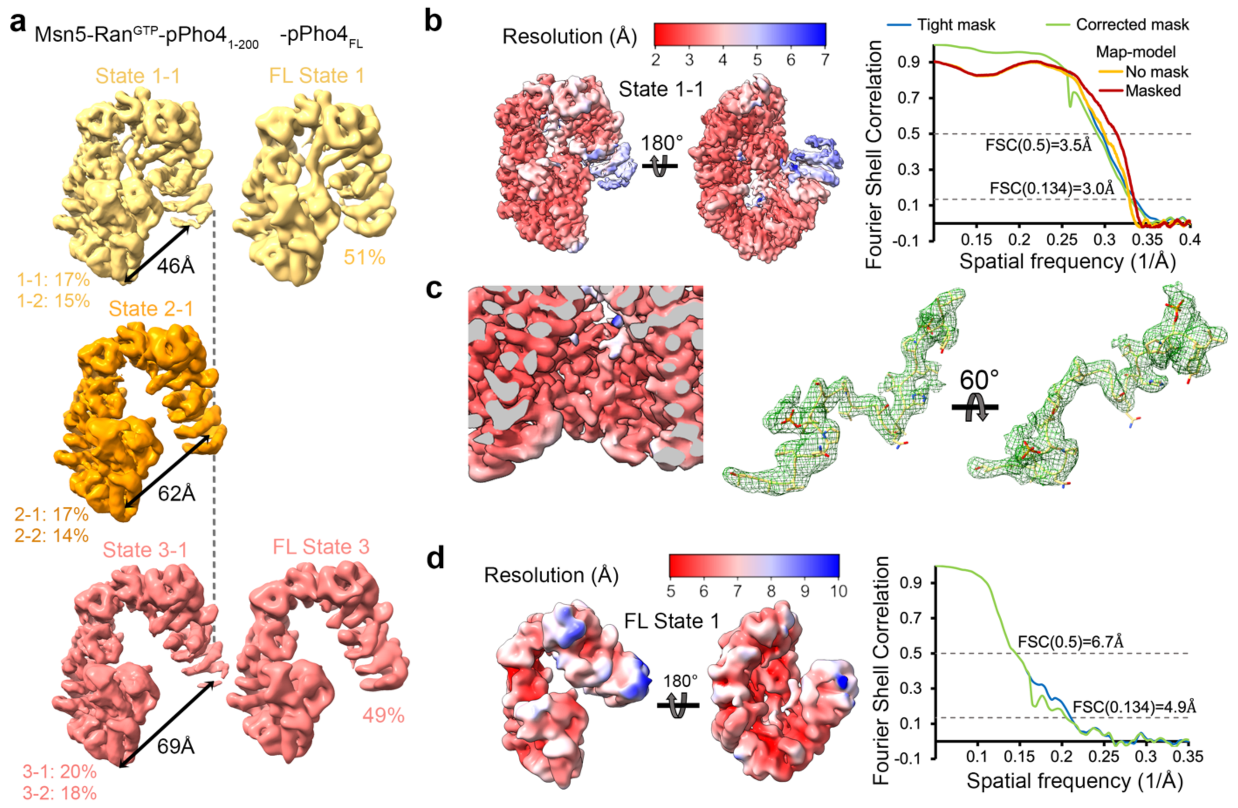


**Extended Data Fig. 2. Different maps of the Msn5-Ran$^{GTP}$-pPho4$_{1\text{-}200}$ and Msn5-Ran$^{GTP}$-pPho4$_{FL}$ complexes. (a)** Left: Particles from the Msn5-Ran$^{GTP}$-pPho4$_{1\text{-}200}$ cryo-EM dataset were separated into 6 classes that are grouped into 3 states based on their Msn5 solenoid conformations. Representative State 1-1 (top, yellow), 2-1 (middle, orange) and 3-1 (bottom, coral) maps are aligned by their Ran$^{GTP}$ regions. Right: Particles from the Msn5-Ran$^{GTP}$-pPho4$_{FL}$ dataset were split into two states. Particle distributions in both the Pho4$_{1\text{-}200}$ and Pho4$_{FL}$ datasets are also indicated. Msn5-Ran$^{GTP}$-pPho4$_{1\text{-}}$particles are about evenly divided across three States: 32% in State 1, which has the most compact Msn5 solenoid, 31% in State 2 and 38% in State 3, which has the most open Msn5 solenoid. Msn5-Ran$^{GTP}$-Pho4$_{FL}$ particles are about evenly distributed into the two states, with 51% in State 1 and 49% in State 3. **(b)** Map statistics and local resolution of the State 1-1 map of the Msn5-Ran$^{GTP}$-Pho4$_{1\text{-}200}$ complex. CryoSPARC FSC curves are plotted. Map-model FSCs are from PHENIX. **(c)** Zoom-in views for the modelled pPho4$_{112\text{-}134}$ peptide of the Msn5-Ran$^{GTP}$-Pho4$_{1\text{-}200}$ structure. **(d)** Shown as in **(b)**, statistics and local resolution of the State 1 map Msn5-Ran$^{GTP}$-Pho4$_{FL}$ complex.

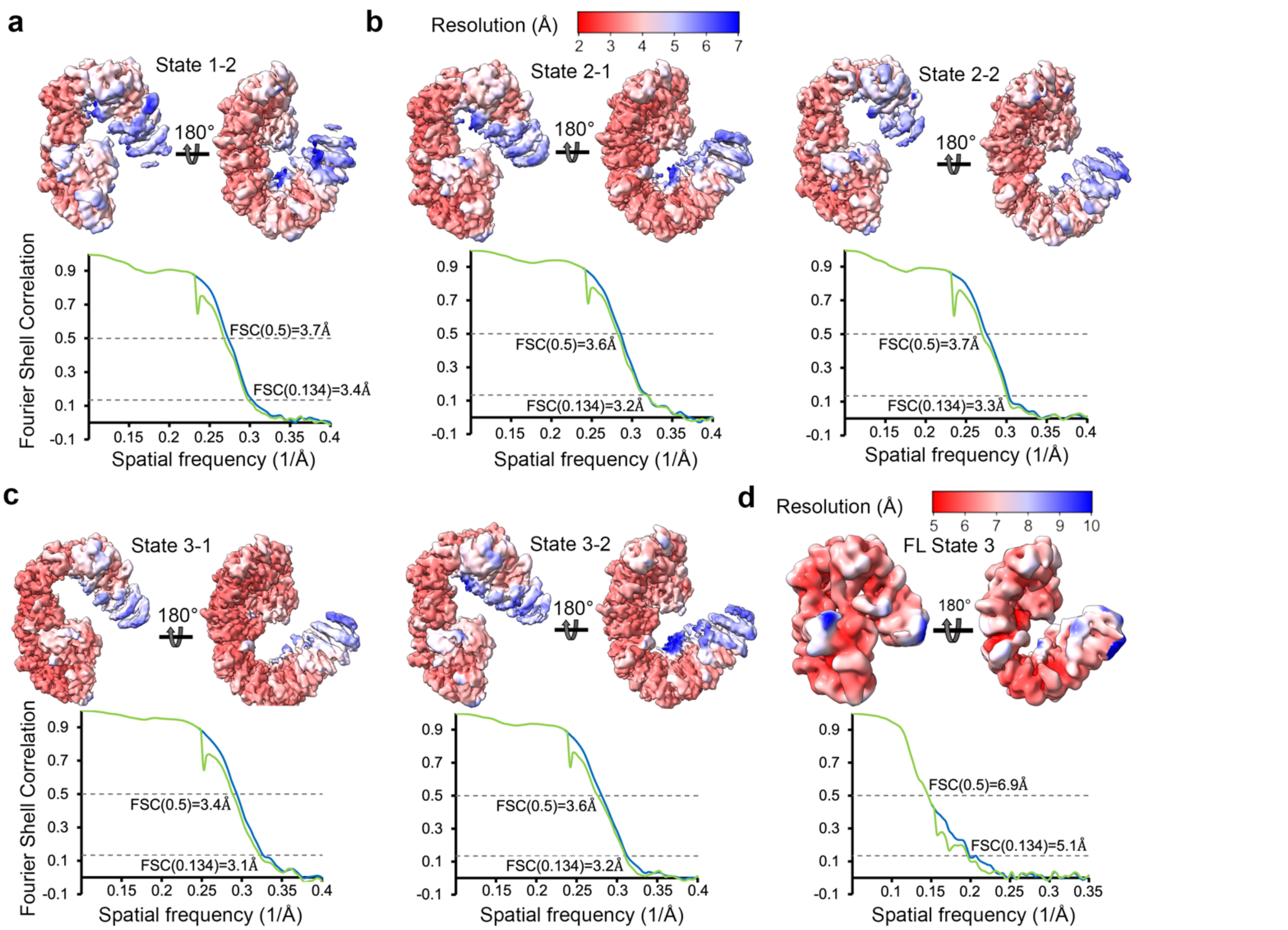


**Extended Data Fig. 3. Statistics of State 1-2, 2-1, 2-2, 3-1 and 3-2 maps of the Msn5-Ran$^{GTP}$-pPho4$_{1\text{-}200}$ complex and State 3 map of the Msn5-Ran$^{GTP}$-pPho4$_{FL}$ complex . (a-c)** Map statistics and local resolution of State 1—2, 2-1, 2-2, 3-1 and 3-2 maps obtained from the Pho4$_{1\text{-}200}$ dataset. cryoSPARC FSC curves are plotted and map-model FSCs are from PHENIX. **(d)** Statistics and local resolution of the State 3 map of the Pho4$_{FL}$ complex.

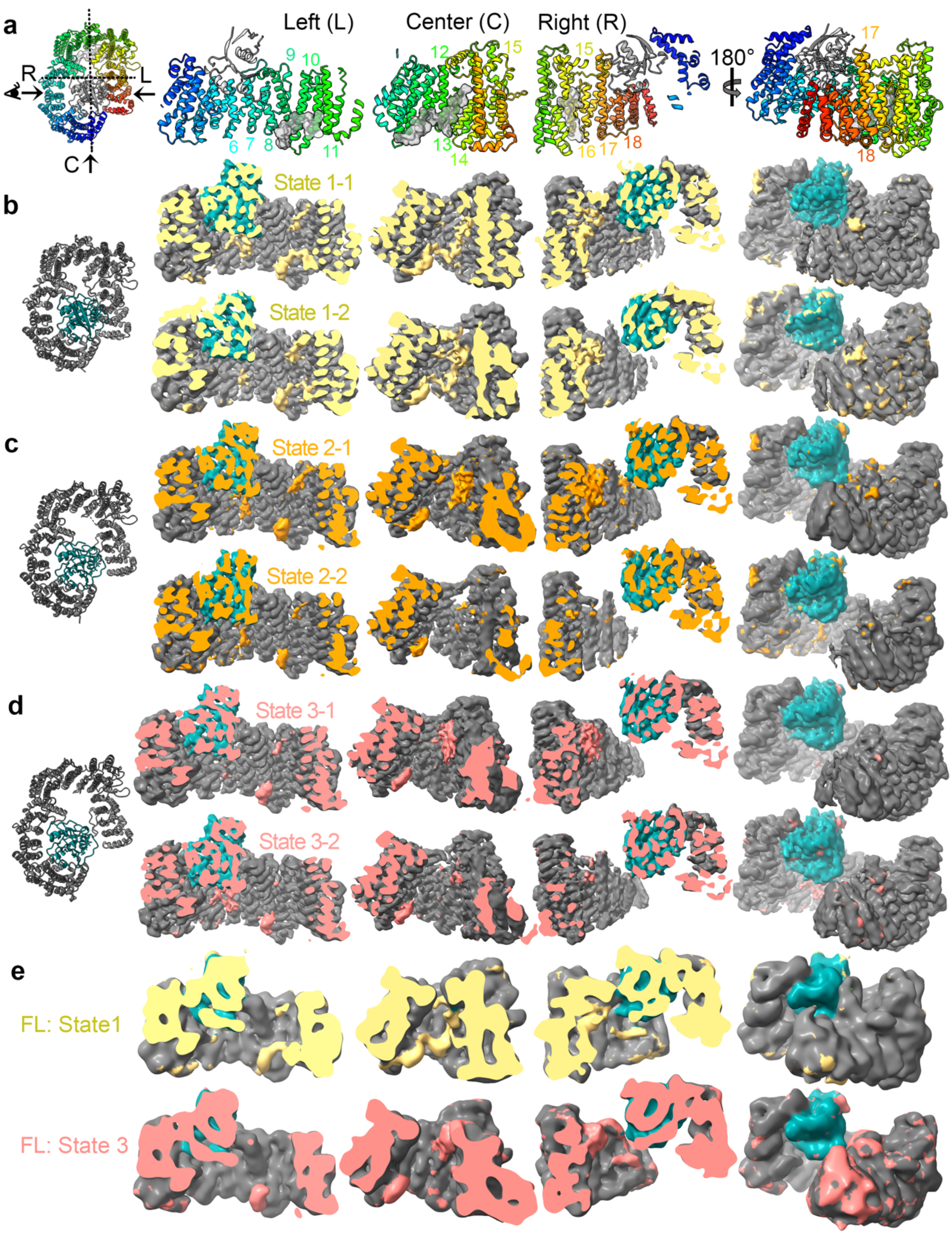


**Extended Data Fig. 4. Cryo-EM maps of Msn5-Ran$^{GTP}$-pPho4 complexes. a)** Cryo-EM structure of Msn5-Ran$^{GTP}$-pPho4$_{1\text{-}200}$ in cartoon (Msn5 colored in rainbow: blue to red = N to C; the rest in grey). View angles for the left (L), right (R) and center (C) are indicated on the top-down view on the left. **(b-e)** Cryo-EM maps of indicated states are colored by the reference model on the left containing only Msn5 (dark gray) and Ran$^{GTP}$ (cyan) so that any extra densities are apparent (yellow, orange and coral for states 1-3 respectively).

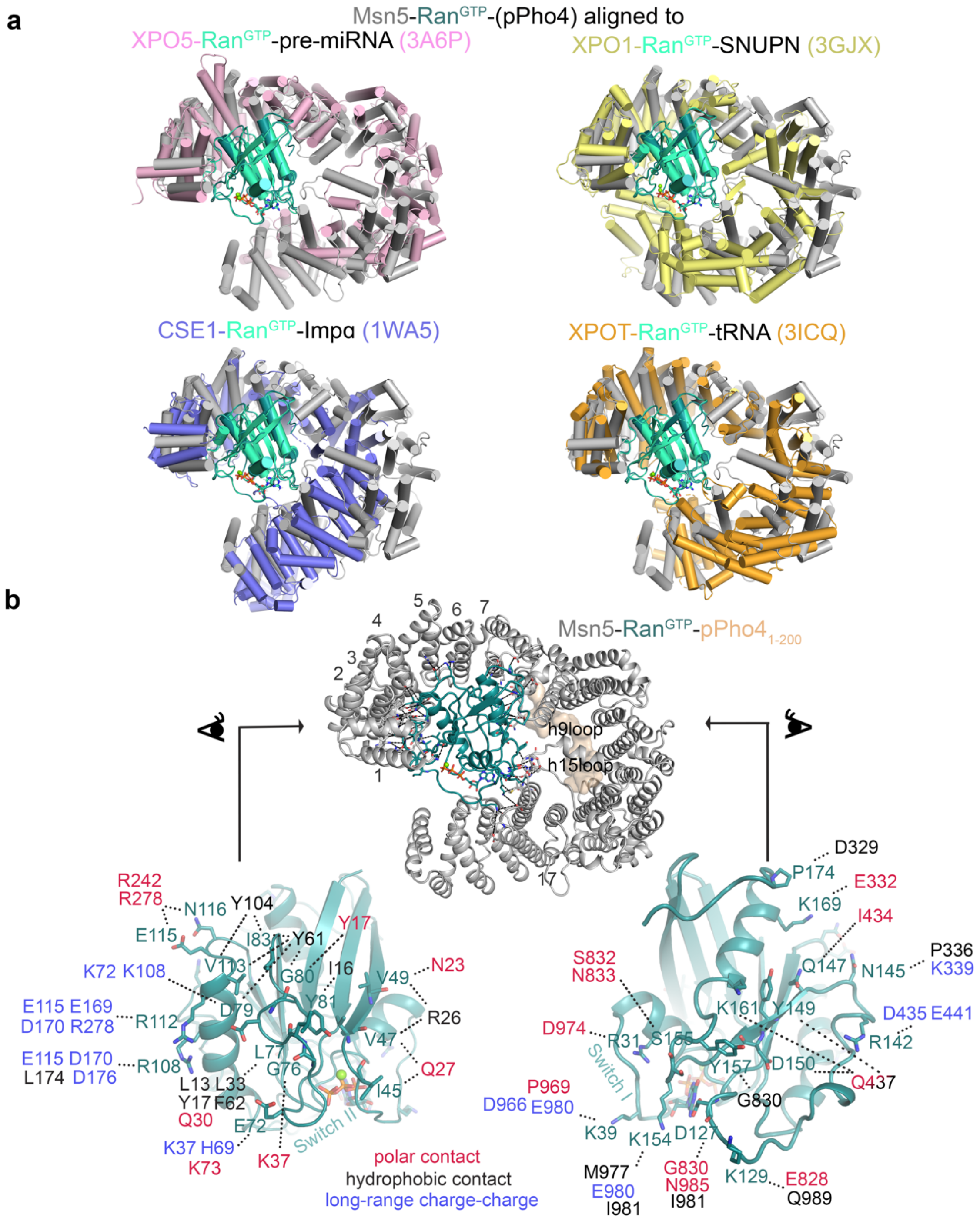


**Extended Data Fig. 5. Interactions between Msn5 and Ran$^{GTP}$. (a)** The cryo-EM structures of Msn5 (gray)-Ran$^{GTP}$ (dark cyan)-pPho4$_{1\text{-}200}$ (not shown) and other Exportin-Ran$^{GTP}$-cargo complexes are aligned by their GTPases ( for clarity, cargoes are not displayed).[9-12] **(b)** Contacts between Msn5 and Ran$^{GTP}$ in Msn5-Ran$^{GTP}$-pPho4$_{1\text{-}200}$ (wheat surface) shown in dotted lines. As there are many contacts that cannot be viewed easily, the front and back views of Ran$^{GTP}$ are displayed below with their interacting residues on Msn5 indicated and colored by the nature of the contact.

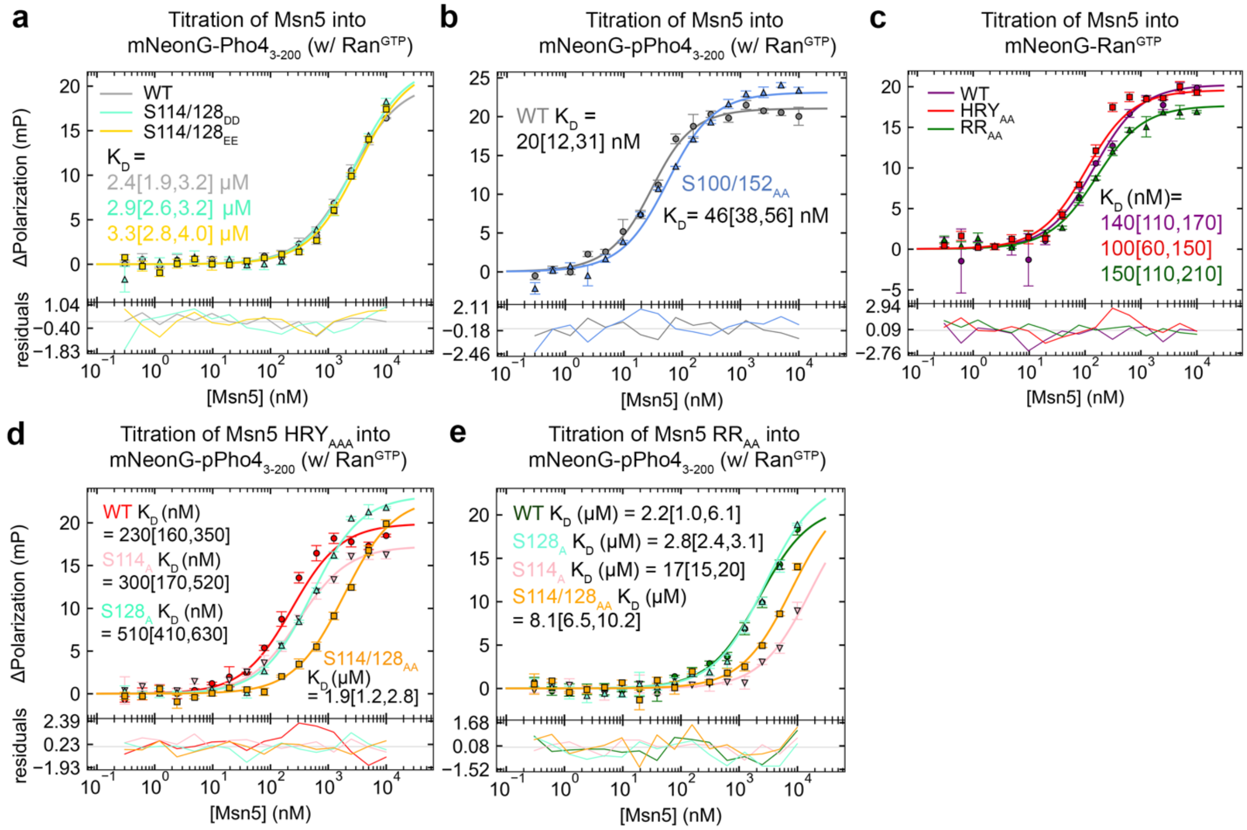


**Extended Data Fig. 6. FP assays of wild-type (WT) and mutant Msn5 binding to phosphorylation site mutants of Pho4. (a)** WT Msn5 binding to unphosphorylated WT (gray), phosphomimic mutant S114/128$_{DD}$ (light cyan; S114 and S128 mutated to aspartic acid) or S114/128$_{EE}$ (light yellow; S114 and S128 mutated to glutamic acid) mNeonG-Pho4$_{3\text{-}200}$, in presence of excess Ran$^{GTP}$. All three mNeonG-Pho4$_{3\text{-}200}$ constructs bind Msn5 with similar affinities. Ineffectiveness of the phosphomimic mutations suggests that the phosphate groups of pS114 and pS128 bind Msn5 via hydrogen-bonding rather than electrostatic interactions. This finding is not entirely surprising as the efficacy of phosphomimic mutations is often case-dependent. **(b)** WT Msn5 binding to Pho4 WT (gray) and S100/152$_{AA}$ mutant (blue; S100 and S152 were mutated to alanine). WT titration is duplicated from **Fig. 2E**. **(C)** Control for WT (purple), HRY$_{AA}$ (red) and RR$_{AA}$ (green) Msn5 from **Fig. 3** binding to mNeonG-Ran$^{GTP}$. **(d-e)** As in **Fig. 3B**, HRY$_{AA}$ (D) or RR$_{AA}$ (E) Msn5 binding to mNeonG-pPho4$_{3\text{-}200}$ S114$_{A}$ (pink), S128$_{A}$ (light cyan) and S114/128$_{AA}$ (orange) mutants. The WT pPho4 traces (red for Msn5 HRY$_{AA}$ and green for Msn5 RR$_{AA}$) are duplicated from **Fig. 3B**. All data points represent mean ± s.d. of triplicate measurements. Lines represent 1-site binding fit and residuals of the fits are plotted below. Dissociation constants ($K_D$) with 95% confidence intervals obtained using error-surface projection are displayed.

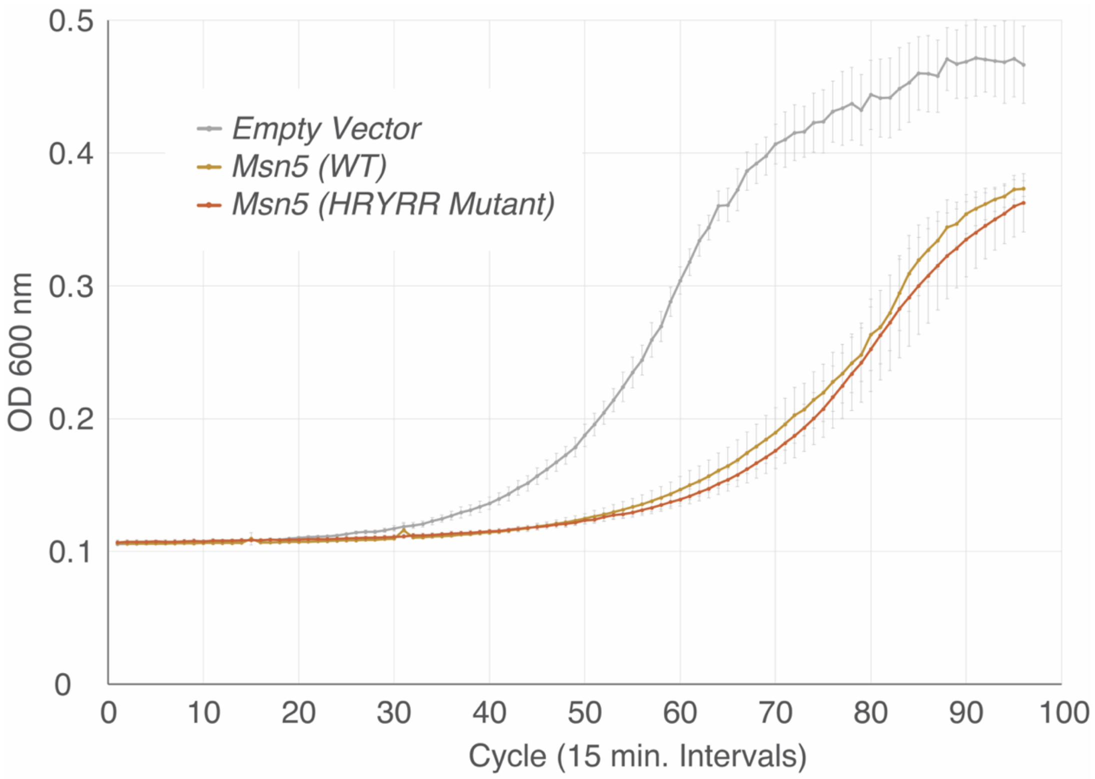


**Extended Data Fig. 7. Growth curve of strains carrying an empty vector control, a WT Msn5 or the HRYRR$_{AAAAA}$ double mutant.** There has been demonstrated evidence that overexpression of Msn5 can cause deleterious effects to cell growth[13-15]. We used the base ASY788 (Msn5 KO) strain and transformed in an empty vector, a WT Msn5, or an HRYRR$_{AAAAA}$ double mutant in a pRS414 vector, with the powerful ADH1 promoter to induce substantial overexpression.

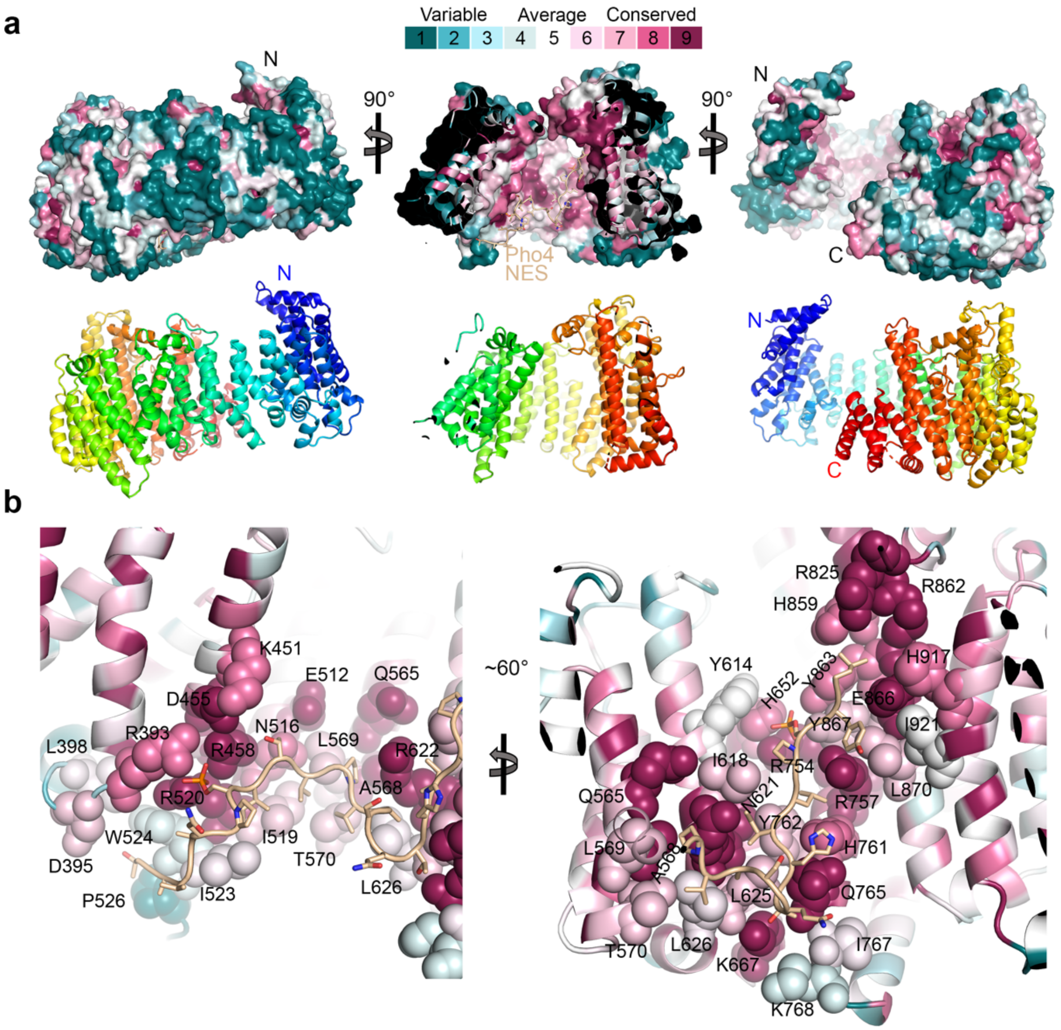


**Extended Data Fig. 8. ConSURF analysis of 101 fungal homologs of Msn5. (a)** Conservation analysis of Msn5 is performed on the ConSURF server[16] and displayed in PyMOL, colored according to the conservation score in the key. Msn5 is shown as surface, and pPho4 NES peptide in wheat cartoon, $Ran^{GTP}$ is not shown. Cartoon of the $Ran^{GTP}$ and $pPho4_{1\text{-}200}$ bound Msn5 in the three views are shown below, colored in rainbow (N to C: blue to red). **(b)** Zoom-in view of the pPho4 NES interface on Msn5. Msn5 residues at the Pho4 interface (by PISA analysis[17]) are drawn as spheres and labelled.

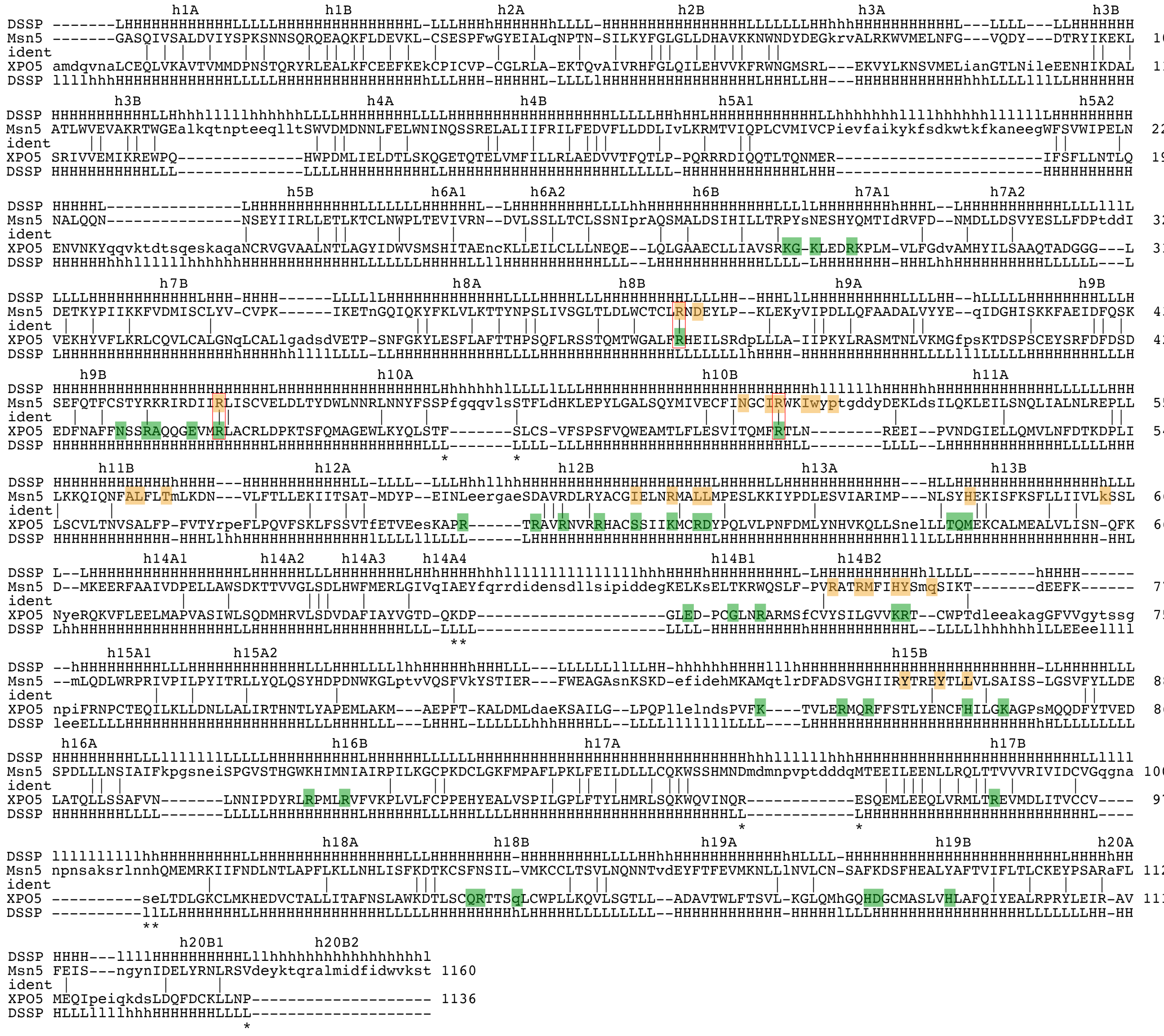

**Extended Data Fig. 9. Alignment of the Msn5 and XPO5 sequences.** Cargo-bound structures of Msn5 and XPO5 (3A6P[9]) were aligned using the DALI server[18]. The aligned residues are in upper case; those that are identical are marked with | and unmodelled sequences not displayed in the alignment are marked with **. Msn5 residues that contact Pho4 are shaded yellow, and XPO5 residues that contact pre-miRNA are shaded green. The conserved arginine residues in Msn5 and XPO5 that contact Pho4 and pre-miRNA, respectively, are marked with red boxes. Msn5 residues that likely interact with Pho4 NES region 1, R996/F1054/N1055, are quite different from the equivalent XPO5 residues (Q1045/R1046) that bind the dsRNA stem.

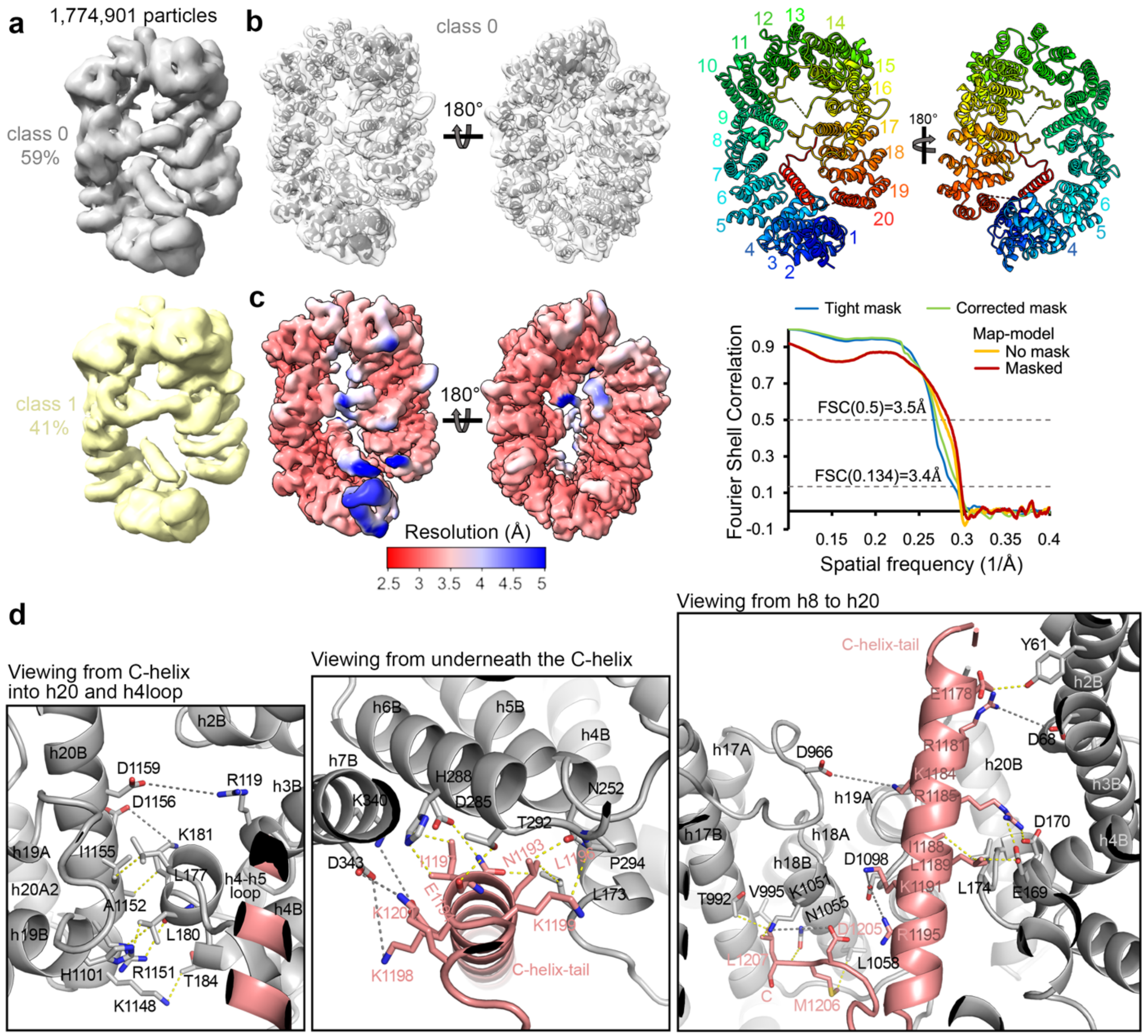


**Extended Data Fig. 10. Unliganded Msn5 cryo-EM maps, model statistics and autoinhibitory contacts. (a)** Distribution of unliganded Msn5 particles in two similar classes obtained in cryoSPARC heterogenous refinement. ~60% of the particles (class 0) were used for reconstruction and structure determination. Reconstruction with the other ~40% of particles (class 1) gave very similar results, but inclusion of these particles did not improve the class 0 map, suggesting small conformational variability/flexibility of unliganded Msn5. **(b)** The final class 0 map, obtained using non-uniform refinement, overlayed with the final structure of unliganded Msn5. The structure is also shown on the right, colored in rainbow, with their HEAT repeats indicated. **(c)** The map in **(b)** colored by local resolution on the left and FSC curves from cryoSPARC (tight and corrected mask; blue and green) and PHENIX map-model FSC (with and without mask; yellow and red) are plotted on the right. **(d)** Msn5 is autoinhibited by intramolecular contacts between the N- and C-terminal HEAT repeats and by two loops, h15loop and h17loop, which reach across the ring to interact with repeats h9-h11. Contacts (<4 Å in yellow and long-range electrostatics of <8 Å in gray) between Msn5 h19-h20 and h3-h4 are shown on the left; between the C-helix-tail (in salmon) with the rest of Msn5 (in gray) in two views on the right.

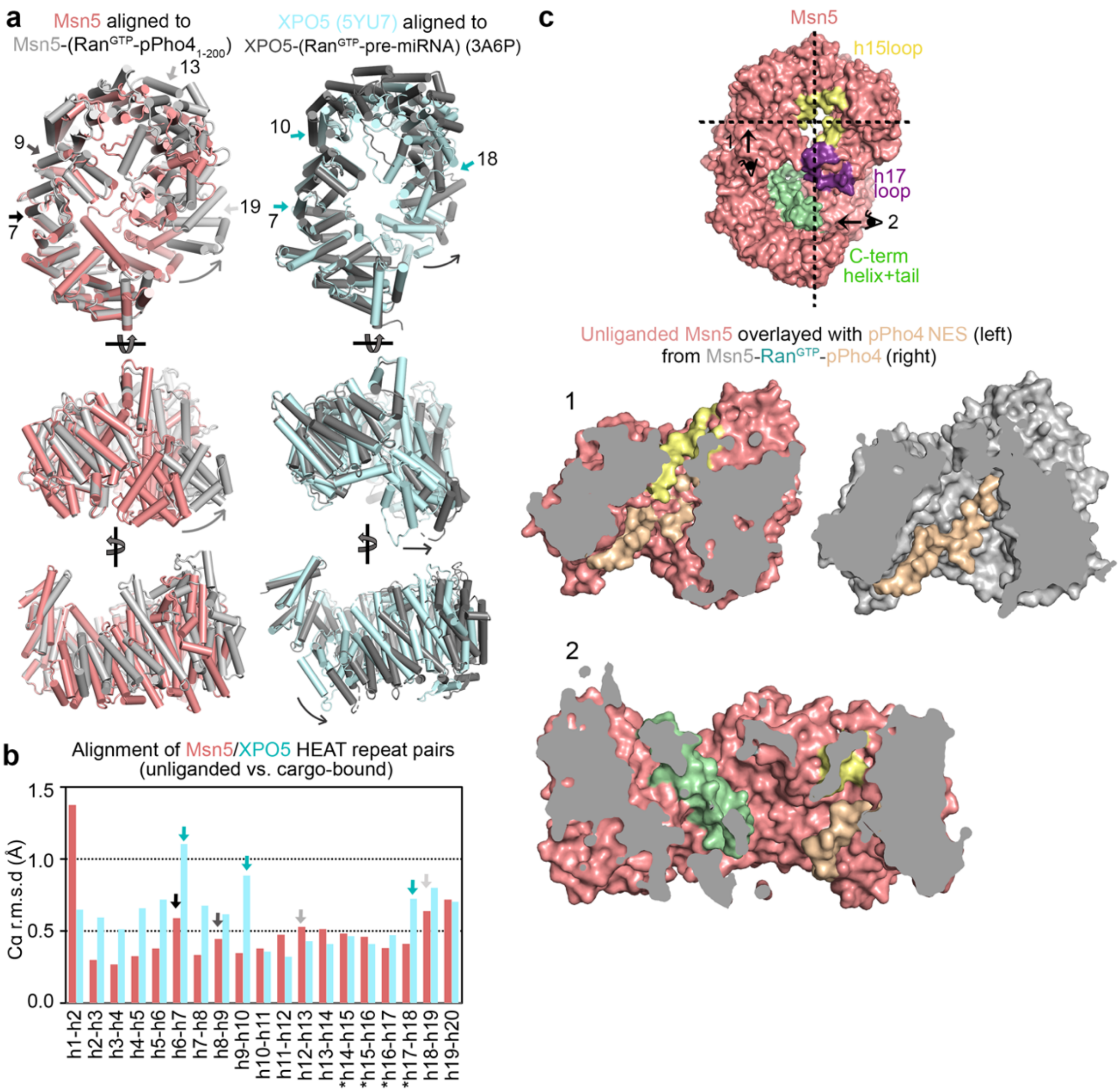


**Extended Data Fig. 11. Comparison of unliganded and cargo-bound Msn5/XPO5 structures. (a)** Left: Unliganded Msn5 unliganded (pink) aligned with Ran$^{GTP}$-pPho4$_{1\text{-}200}$-bound Msn5 (light grey; Ran and Pho4 not shown). Right: Unliganded XPO5 unliganded (cyan; 5YU7) aligned with Ran$^{GTP}$-premiRNA-bound XPO5 (dark grey; Ran and pre-miRNA not shown; 3A6P).[9,19] Both Msn5 and XPO5 solenoids are more open when bound to cargo. **(b)** C$\alpha$ root-mean-square-deviation (r.m.s.d) of aligned HEAT repeat pairs of unliganded and cargo-bound Msn5 (pink) and XPO5 (cyan) structures shown in **(a)**. No large changes are observed at any HEAT repeats. Structural differences are distributed across the whole solenoid with small increases in r.m.s.d at a few HEAT repeats that are marked with arrows (also marked in **(a)**) that may act as hinges. **(c)** Unliganded Msn5 structure shown as colored surface in two cut-out views, view 1 (middle) and view 2 (bottom), as indicated on the top-down view on the top. In left panel for view 1, pPho4 NES overlayed from the Msn5 (gray)-Ran$^{GTP}$ (not shown)-pPho4$_{1\text{-}200}$ (wheat) structure (right panel) clash with the unliganded Msn5 – compare visibility of the pPho4 NES in right and left panels. In view 2, the entire pPho4 NES region 4 can be seen with no obstruction from unliganded Msn5.

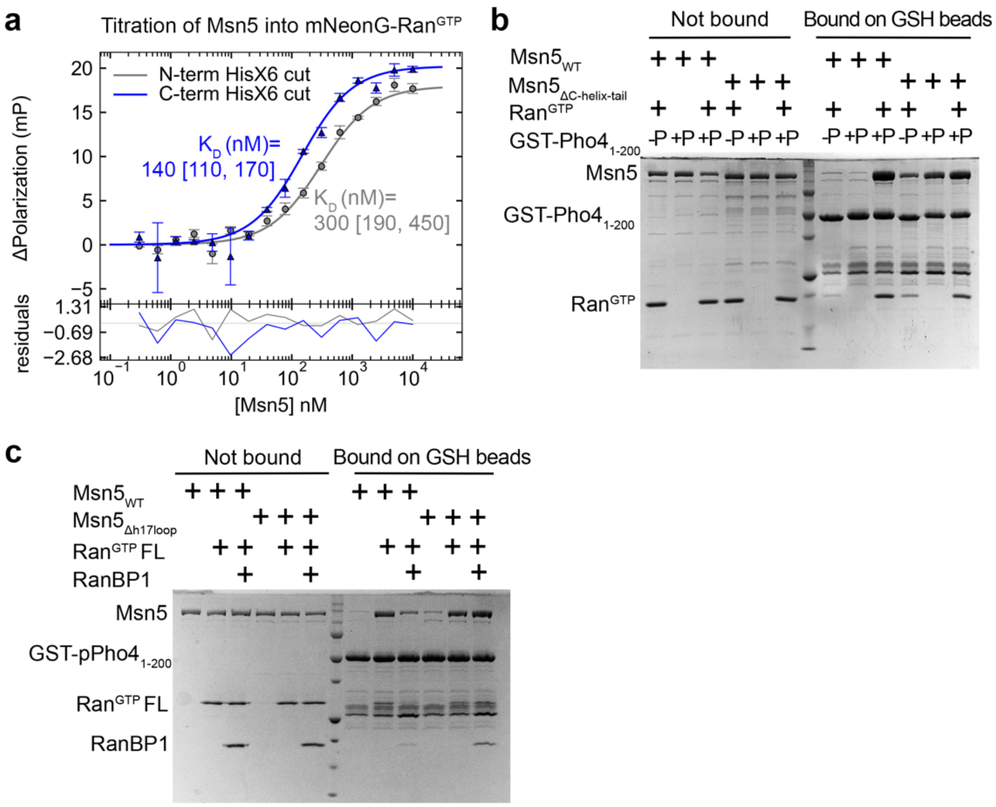


**Extended Data Fig. 12. Binding assays of Msn5 with Ran and RanBP1. (a)** Cryo-EM structure determination was performed using Msn5-HisX6 where the HisX6 tag was removed by Tev protease. Here, FP analysis was performed with HisX6-Msn5 and Msn5-HisX6 where the N- or C-terminal HisX6 tags, respectively, were removed with TEV protease. Msn5 was titrated into 20 nM mNeonG-$Ran^{GTP}$ and polarization signal was measured. Data points represent mean ± s.d. of triplicate measurements. Line represents 1-site binding fit and residuals of the fits are plotted below. Dissociation constants ($K_D$) with 95% confidence intervals is displayed. Msn5 binds $Ran^{GTP}$ with $K_D$ 140 nM or 300 nM. Such moderate affinity of Msn5 for $Ran^{GTP}$ in the absence of cargo is unusual exportins, which usually bind with much lower affinities to $Ran^{GTP}$ alone.[20] This finding suggests that although a large population of unliganded Msn5 adopts a compact ring conformation, the autoinhibited ring has some flexibility and small populations of other states with open rings and exposed $Ran^{GTP}$ binding site also exist. The presence of additional residues from the TEV cleavage site on the C-terminus of Msn5 increases $Ran^{GTP}$ affinity 2-fold, supporting the importance of the Msn5 C-terminus in stabilizing the closed-ring autoinhibited Msn5 conformation. **(b-c)** Full gels for the binding assays in **Fig. 4C** and **E**, which include the unbound proteins of the assays.

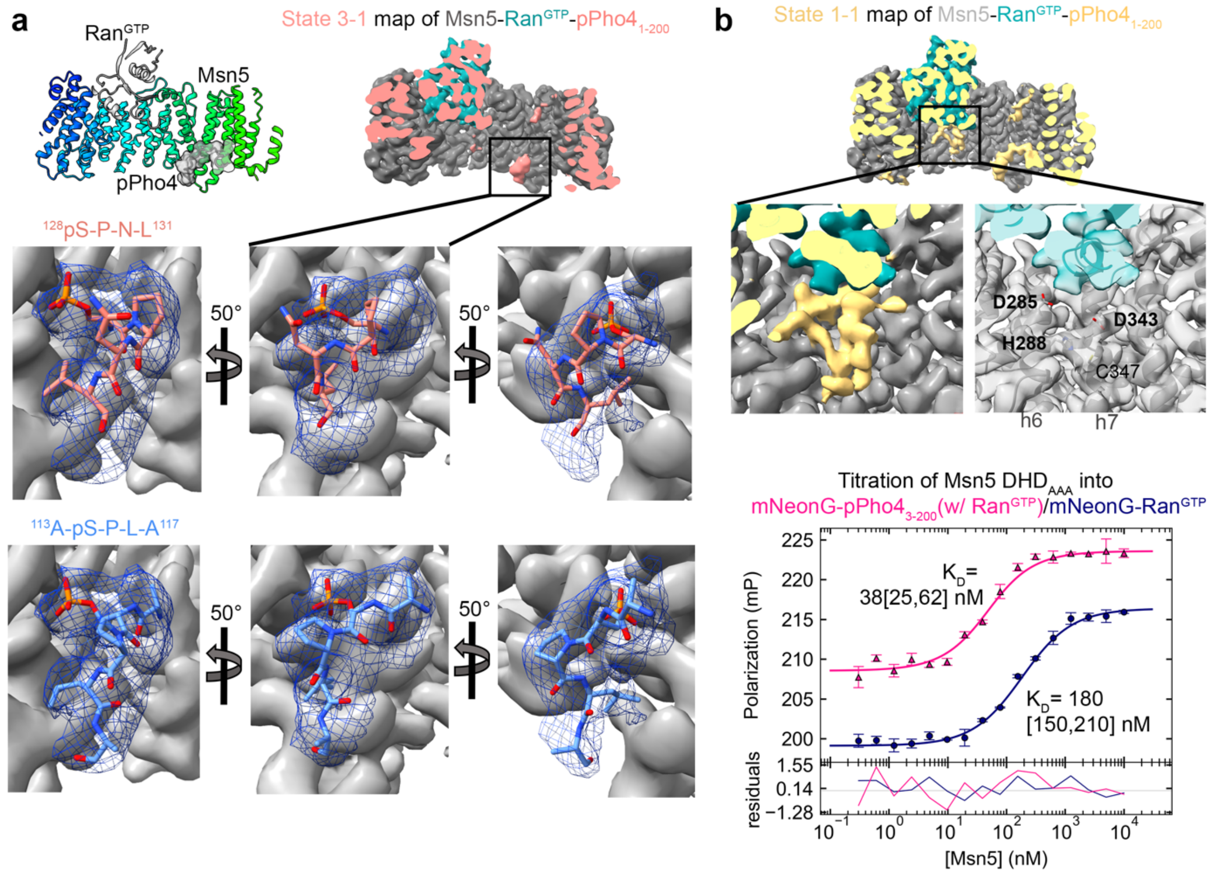


**Extended Data Fig. 13. CryoEM map densities that cannot be confidently modeled, see Supplementary text. (a)** The Msn5 (rainbow cartoon)-Ran$^{GTP}$(gray cartoon)-pPho4 (gray cartoon and surface) structure is shown on top left as in **Fig. S3A**. The Pho4 density in State 3-1 map of Msn5 (gray)-Ran$^{GTP}$ (cyan)-pPho4$_{1\text{-}200}$ (salmon) is shown on top right, marked by a box and in the same orientation as the structure on the left. Six zoom-in panels of the boxed area are shown below with Msn5 density in grey and the Pho4 density near the Msn5 RR site as blue mesh. The center 2 panels are in approximately the same orientation as the boxed area. Two possible pPho4 models, each with a phosphoserine ($^{128}$pSPNL$^{131}$ in salmon or $^{113}$ApSPLA$^{117}$ in blue), can be fitted into the Pho4 density. The $^{113}$ApSPLA$^{117}$ model (blue) seems to fit better in all orientations. However, it is unclear which one is correct, and it is also possible that the density represents a mix of both models. **(b)** The state 1-1 map of the Msn5 (gray)-Ran$^{GTP}$ (cyan)-pPho4$_{1\text{-}200}$ (yellow) complex with unmodeled and unexplained density close to Ran$^{GTP}$ marked with a box (top). Middle left panel: Zoom-in view of unexplained density. Middle right panel: The same map is shown transparent, to see the Msn5 cartoon and sticks below the transparent extra density. Bottom panel: FP titrations of Msn5 mutant where D285/H288/D343 are mutated to alanines (DHD$_{AAA}$) binding to mNeonG-pPho4$_{3\text{-}200}$ in the presence of excess Ran$^{GTP}$ (pink) or to mNeonG-Ran$^{GTP}$ (navy). Data points represent mean ± s.d. of triplicate measurements. Line represents 1-site binding fit and residuals of the fits are plotted below. Dissociation constants ($K_D$) with 95% confidence intervals obtained using error-surface projection are displayed.
